# Heterogeneity in neural network engagement supports individual differences in top-down attention

**DOI:** 10.64898/2026.01.27.701998

**Authors:** Emma Holmes, Karl J. Friston, Timothy D Griffiths

## Abstract

Top-down attention enables selective prioritisation of goal-relevant information, yet how attentional control varies across individuals remains unclear, particularly in relation to sensory processing. Here, we combined magnetoencephalography (MEG) and pupillometry—in older adults performing a spatial cueing task—to examine both shared and subject-specific aspects of top-down attention. Informative spatial cues elicited sustained activity in fronto-parietal and sensory regions, alpha lateralisation, and increased pupil dilation, replicating established neural and physiological markers. Crucially, these group-level patterns were accompanied by systematic intersubject variability: individuals with greater hearing loss showed increased preparatory engagement of precentral and postcentral gyri, and variation in pupil dilation predicted activity in frontal regions. These results reveal that top-down attention arises from heterogeneous network configurations shaped by sensory processing and strategic resource allocation. Overall, our findings underscore the need to move beyond group averages to capture the neural architecture underlying individual differences in top-down attention.

## Introduction

Top-down (i.e., ‘endogenous’) attention enables humans to selectively prioritise goal-relevant information in complex environments. The functional architecture of top-down attention is well-established at the group level, consistently engaging fronto-parietal areas^1,2^ that include the intraparietal sulus (IPS), superior parietal lobule (SPL), inferior frontal gyrus (IFG), and dorsal premotor and prefrontal areas (e.g., homologues of frontal eye fields), with feedback modulating responses in relevant sensory regions^3,4^. Yet, most previous research assumes that individuals engage similar mechanisms and few studies have examined the neural bases of individual differences in top-down attention in non-clinical populations. Here, we test the hypothesis that top-down attention is not mediated by a single functional architecture, but by heterogenous, subject-specific network configurations that reflect strategic variation across individuals. Characterising strategic variation is crucial for interpreting the implicit cognitive degeneracy^5–7^—in other words, a many-to-one structure-function mapping—in both clinical and non-clinical populations.

Given the intimate interplay between perception and attention, variation in sensory processing could influence how individuals deploy top-down attention. Sensory processing varies substantially across individuals, in both young^8,9^ and ageing^10,11^ populations, and correlates with broader cognitive changes in dementia^12–14^. Notably, even minimal, sub-clinical sensory loss reduces the behavioural benefit that individuals gain from attentional cues^15^, suggesting that variability in sensory precision affects the deployment of top-down attention. Such variability may shape the functional architecture for top-down attention, which would be consistent with functional reorganisation in studies examining more extreme sensory losses^16,17^.

Individual differences in sensory processing could shape top-down attention in at least two distinct ways. First, sensory loss may weaken engagement of canonical attentional networks, particularly when sensory processing reduces the fidelity of target representations. From a predictive coding perspective, this could reflect diminished precision-weighting of sensory signals^18^. From an alternative view, resource-based accounts suggest that greater perceptual demands deplete cognitive reserve^19,20^, which could reduce the flexibility of attentional control. Consistent with these accounts, older adults who experience age-related declines in sensory processing show reduced behavioural benefits of attentional cues^15^ and weaker lateralised alpha power during spatial attention^21,22^. Alternatively, poorer sensory acuity may trigger compensatory recruitment. This could include greater recruitment of canonical networks, and recruitment of additional brain regions. In particular, older adults more readily engage cingulo-opercular regions (including IFG, dorsal cingulate cortex, and anterior insula) during speech perception^23–25^, and such recruitment has been linked to improved speech comprehension^24–29^. Cingulo-opercular regions could facilitate perception through functional integration with fronto-parietal attention regions^30–33^ or because they are associated with allocating mental effort to overcome challenge^34^. Disentangling whether reduced engagement, compensatory recruitment, or a combination of mechanisms underlie heterogeneity is crucial for identifying the mechanisms through which intersubject sensory variation gives rise to individual differences in attention.

Studying pupil dilation is interesting in the context of these accounts. Pupil dilation is a putative index of effort^35,36^, scaling with activity in the locus coeruleus noradrenergic system^37,38^. The locus coeruleus is closely coupled with cingulo-opercular regions^39^ and may drive cholinergic activity relevant to attentional neuronal modulation^40,41^. One line of research relates pupil dilation to the ‘intensity’ of attention (i.e., how much attention is allocated)^42^. Both older adults and young adults with hearing loss show greater pupil dilation during challenging listening^43^, suggesting increased effort to sustain comprehension. From these studies, it is unclear whether greater pupil dilation reflects additional deployment of top-down attention or changes in processing of the target under reduced sensory acuity. Nevertheless, pupillometry offers a way to test heterogeneity in strategic allocation of resources in paradigms that separate top-down attention from target processing.

Here, we provide a novel perspective on variation in top-down attention by simultaneously measuring functional brain activity and pupil responses, and by relating these responses to individual differences in sensory processing. Crucially, we used a paradigm^2,15,44,45^ that allowed us to separate the processes of attentional control from sensory processing of the target. We demonstrate that the functional architecture for top-down attention varies systematically with sensory acuity, and reveal additional individual differences related to the strategic allocation of cognitive resources that map onto pupil dilation. Overall, our findings demonstrate that top-down attention is implemented through degenerate neural network engagement that varies across individuals.

## Results

To study individual differences in top-down attention, we used a modified ‘cocktail party’ listening task^46^, in which participants heard three spoken phrases—each including a colour and number word— from different simulated locations (left, centre, and right), and received a visual spatial cue indicating the target location (i.e., the source of the spoken phrase to be identified; Fig. 1). Crucially, we included a visual pre-cue, presented 2000 ms before the target stimuli, that was either informative about the target location (with 100% validity) or was uninformative. Thus, responses to the cue reflect preparatory top-down attentional control without any confounding sensory processing of the target^2,3,47,48^. This task is well-suited to probing individual differences: young adults with clinically normal hearing show behavioural benefits from advance cueing^49–51^, and this benefit is preserved at the group level in older adults^15^, but decreases with hearing loss^15,52,53^. Moreover, individuals with normal hearing show sustained cue-related responses^45,54^ and alpha lateralisation^51,55–57^ at the group level, but these responses are weaker in children with early-onset hearing loss^45^ and in older adults with suspected age-related hearing loss^21,22^. Unlike other studies examining individual differences during speech perception^58–61^, this task allows us to probe heterogeneity in attentional control, independent of sensory encoding.

**Fig. 1:**
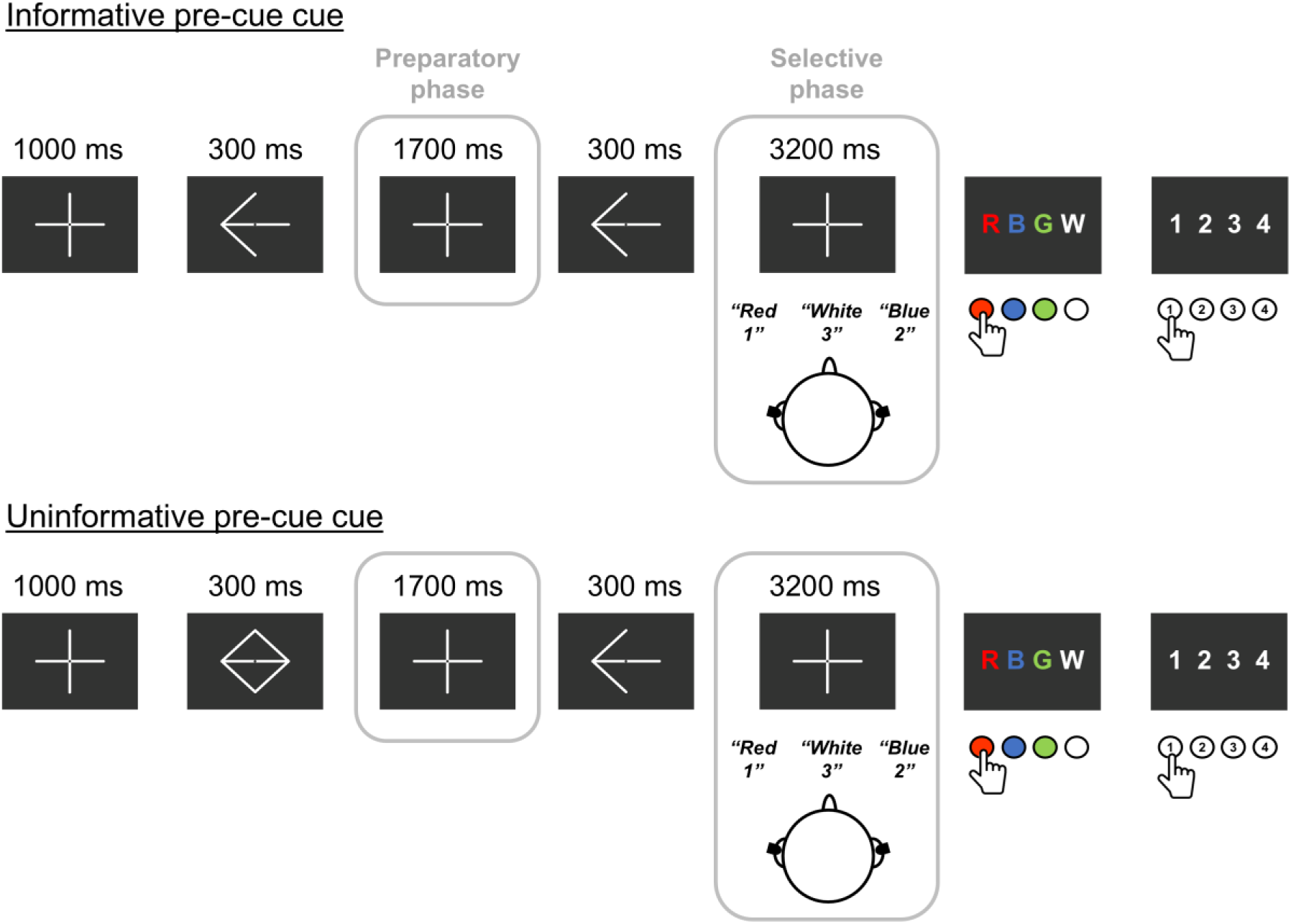
A spatial attention task that isolates top-down attention from target processing. Schematic of the spatial attention task for attend-left conditions. The pre-cue was either a preview of the leftwards-pointing arrow (informative pre-cue condition) or a visual composite stimulus (uninformative pre-cue condition). During the selective phase, stimuli were equivalent in the informative and uninformative pre-cue conditions. The attend-right conditions were equivalent to those shown, except all arrow stimuli pointed rightwards.

We recorded neuronal responses using magnetoencephalography (MEG) in 44 older adults (56–80 years), with simultaneous pupillometry in a subset of 30 participants. As an index of auditory sensory processing, we measured high-frequency (4–8 kHz) audiometric thresholds in each participant, which are sensitive to age-related hearing loss^11^. To facilitate comparisons across participants with different levels of hearing loss, we presented the stimuli at a fixed level above each participant’s 50% threshold for reporting individual spoken phrases.

### Group-level behavior and neural signatures of attention

As an initial step, we examined behavioral performance to confirm that participants engaged with the task as intended. Across the group, participants were more accurate at reporting the target phrase when the pre-cues were informative compared to uninformative (*F*(1, 43) = 5.33, *p* = .026, *ω_p_^2^* = .00 [95% CI = .00, .10]), replicating the established behavioral benefit of advance cueing. As expected, there was no evidence that performance varied according to the direction of attention (main effect of attend-left versus attend-right trials: *F*(1, 43) = 3.19, *p* = .08, *ω_p_^2^* = .00 [95% CI = .00, .11]; interaction between pre-cue and direction: *F*(1, 43) = 1.91, *p* = .17, *ω_p_^2^* = .00 [95% CI = .00, .06]).

We next examined group MEG responses at the sensor level, to test for sustained cue-related activity and alpha lateralisation. A cluster-based permutation analysis, across all sensors, showed greater responses in the informative condition (one significant cluster: time window - 1839.2 to −35.0 ms; sum of *t*-values = 5965.30, *p* = .005), reflecting a sustained difference between informative and uninformative trials during the preparatory phase (Fig. 2a). This effect mirrors responses previously observed in young participants with clinically normal hearing^45,54^. Comparable effects were present when trials were subdivided into attend-left and attend-right trials (Supplementary Fig. 1a), but no such differences were present in a passive visual control task in which participants only viewed the visual stimuli (Supplementary Fig. 1b). Scalp maps indicated lateralised alpha activity during the selective phase, and to a lesser extent during the preparatory phase, when comparing attend-left and attend-right cue conditions (Supplementary Fig. 1c).

**Fig. 2:**
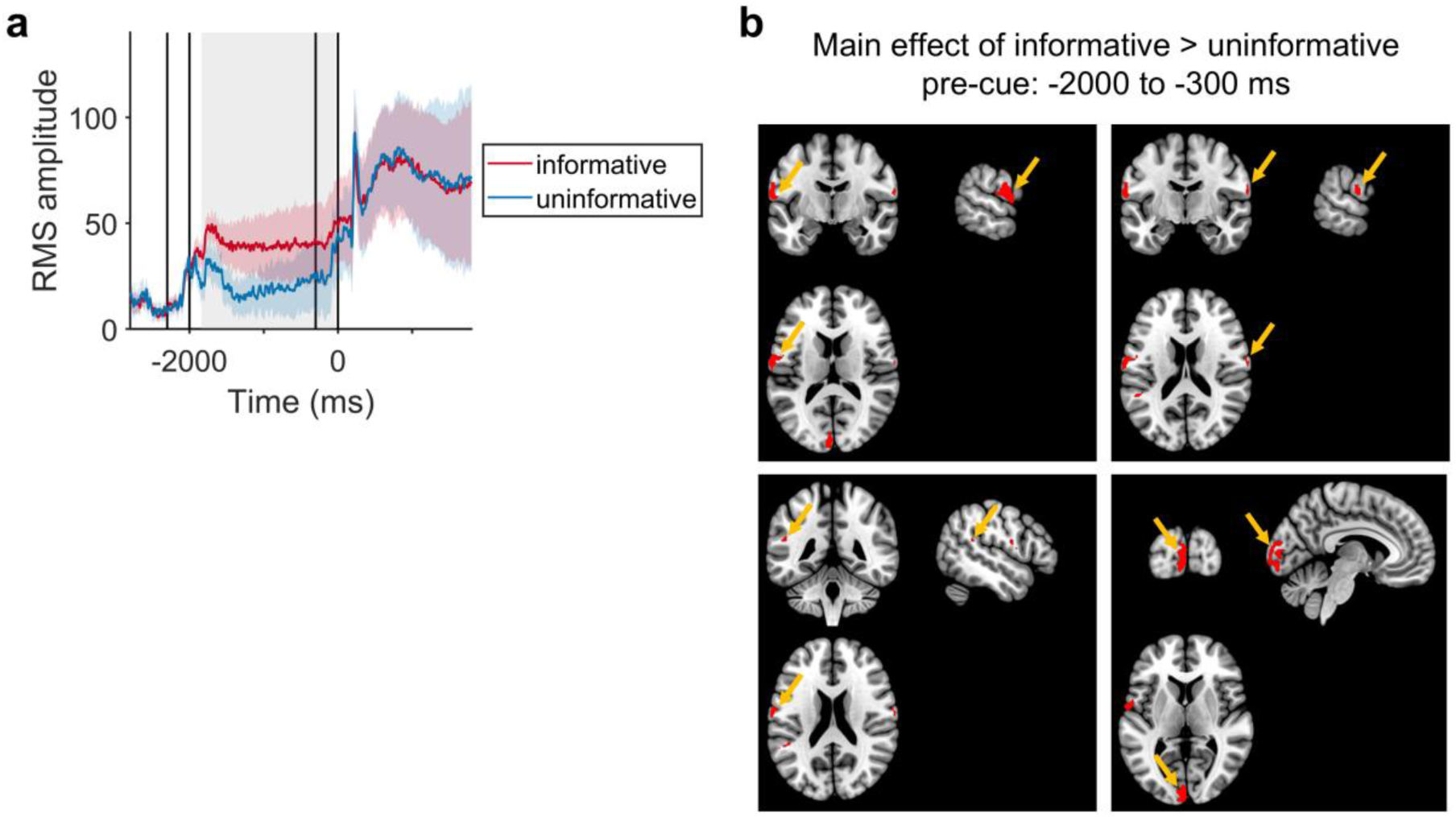
Group-level results. **a**, Magnetic evoked fields across all participants (N = 44) in the informative and uninformative conditions. Lines show the means and standard errors. The shaded box indicates the significant cluster window. **b**, Source analysis during the preparatory phase, showing the difference between the informative and uninformative conditions, at the group level.

Finally, we examined the cortical sources of group-level effects. Using a general linear model, we compared the informative and uninformative conditions during the preparatory phase (−2000 to −300 ms; Fig. 2b). In the informative condition, we observed greater activity in visual cortex, bilateral inferior parietal lobule, and bilateral auditory cortex, extending into the operculum and postcentral gyri (Supplementary Table 1). Inspecting 100-ms sub-windows within the preparatory interval, the most consistent responses across the group emerged during the middle of the preparatory phase (Supplementary Fig. 2; Supplementary Table 2). This activity included the regions identified across the entire preparatory interval, as well as the left superior parietal lobule, bilateral IFG, and bilateral insula. Thus, consistent with prior work^1–4^, informative cues engaged fronto-parietal regions and bilateral auditory cortex, which likely reflects top-down biasing of areas for target processing, given the absence of acoustic input during this interval. We also observed responses in visual cortex, which may reflect maintenance of the informative visual cue in working memory^3,62^.

When comparing how informative and uninformative cues affected processing of the acoustic stimuli during the selective phase, we found greater activity for the informative condition in left auditory cortex (Supplementary Fig. 1d; Supplementary Table 4). This result demonstrates that preparation modulates subsequent responses to acoustic stimuli, potentially reflecting enhanced processing when participants had longer to prepare for the target location.

A source analysis during the passive visual control task showed no significant differences in responses to the informative and uninformative cue stimuli, implying that the effects we observed in the spatial attention task reflect attentional processes rather than responses to the visual stimuli.

Together, these findings validate the task as an effective probe of attentional control, providing context for subsequent analyses of individual differences.

### Individual differences related to sensory processing

To characterise individual differences, we entered between-subject covariates into the above source analyses. We controlled for age (56–80 years), the level of acoustic stimulation determined by each participant’s speech reception threshold (SRT; Fig. 3a), and overall performance on the spatial attention task (‘Accuracy’; Fig. 3b), by entering these three variables as covariates into the general linear model. To examine how differences between informative and uninformative conditions related to hearing acuity, we also entered audiometric thresholds (averaged across the ears and across 4–8 kHz frequencies; ‘Audiogram’; Fig 3c) as an additional effect. These subject-specific effects were used to explain (informative versus uninformative) differences in MEG responses and therefore model interactions with pre-cue effects.

**Fig. 3.**
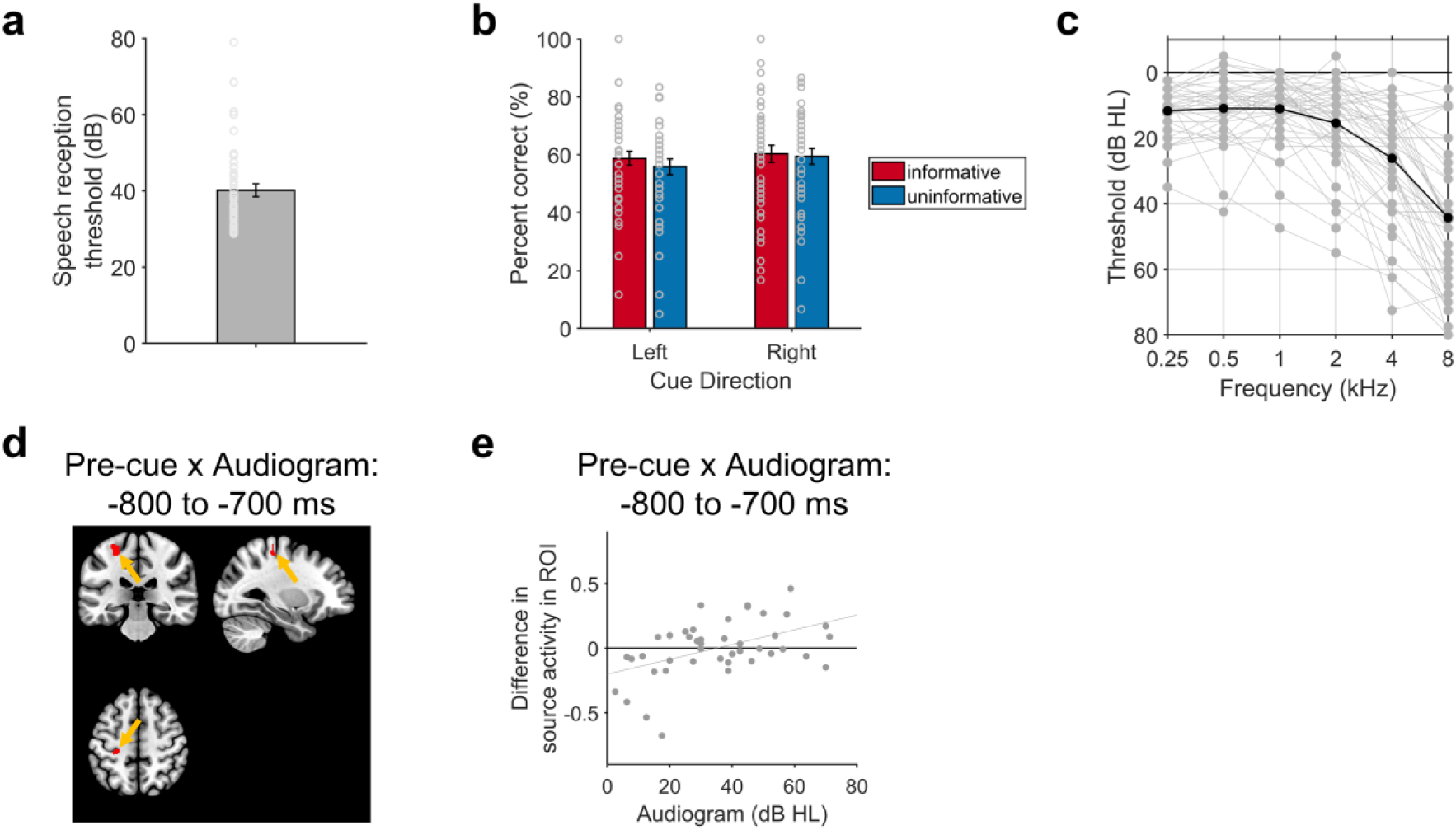
Individual differences among participants. **a**, Speech reception thresholds (mean, standard error, and results from individual participants). **b**, Accuracy in the spatial attention task, separated by pre-cue conditions and into attend-left and attend-right trial types (each bar displays the mean, standard error, and results from individual participants). **c**, Pure-tone audiometric thresholds at .25–8 kHz, recorded in decibels hearing level (dB HL), and averaged across the left and right ears. The solid black line shows mean thresholds across the group, and grey lines show thresholds for individual participants. **d**, Results of the source localisation in the −800 to - 700 ms time window, showing the interaction between the Audiogram (at 4–8 kHz) and the effect of the pre-cue. **e**, Scatter plots showing the relationship in d (grey circles show data points from each participant, and grey line shows the line of best fit).

When considering the entire preparatory phase (−2000 to −300 ms), none of the covariates showed significant effects. Also, no effects of age, SRTs, or performance emerged in the sub-window analysis. However, audiometric thresholds showed effects at two time windows (Supplementary Table 3). These time windows (−1400 to −1300 ms, and −800 to −700 ms) were during the middle of the preparatory phase. Hearing acuity was associated with activity in the left precentral and postcentral gyri in the informative condition relative to the uninformative condition (Fig. 3d). Specifically, individuals with more age-related hearing loss showed greater activity in this source (Fig. 3e), suggesting that individuals who have greater hearing loss engage preparatory activity in the left precentral and postcentral gyri when they have advance knowledge of target location.

During the selective phase, we found no significant interactions between any of the covariates and the informative and uninformative effect.

### Relationships between preparatory spatial attention and pupil dilation

We next examined pupil dilation in a subset of 30 participants who had usable pupillometry data. Compared to MEG responses, pupil dilation evolves over a slower timescale^63^.

We found that pupil dilation was greater in the informative than in the uninformative condition during the preparatory phase (Fig. 4a; one significant cluster: time window −1546 to −374 ms; sum of *t*-values = 2225.5, *p* = .028).

**Fig. 4.**
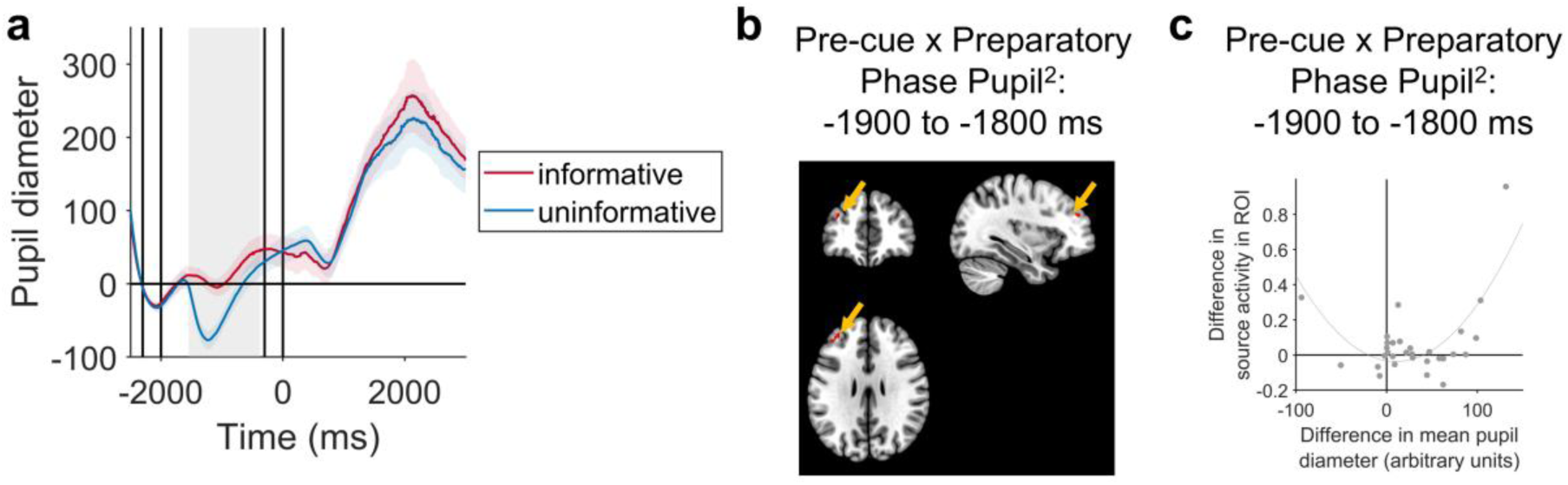
Pupil dilation results. **a**, Pupil diameter in the informative and uninformative conditions, across participants who had usable pupil data (N = 30). Lines show the means and standard errors. The shaded box indicates the significant cluster window. **b**, Results of the source localisation in the - 1900 to −1800 ms time window (indicative of the significant time windows), showing the interaction between the squared pupil dilation during the preparatory phase and the effect of the pre-cue (informative - uninformative). **c**, Scatter plot showing the relationship in b (grey circles show data points from each participant, and grey line shows the line of best fit).

To examine the relationship between MEG and pupil responses, we used a general linear model contrasting the informative and uninformative conditions with four covariates derived from pupil diameter. The first two covariates captured the difference in mean pupil diameter between informative and uninformative trials in the preparatory and selective phases, respectively (with time windows defined *a priori*). The remaining two covariates represented the squared differences in mean pupil diameter for each phase, to account for potential non-linear (e.g., Yerkes-Dodson) associations with neural activity^64,65^.

As in the earlier covariate analyses, we found significant effects in the sub-window analysis but no significant effects across the entire preparatory interval (−2000 to −300 ms). During the first half of the preparatory phase (in time windows between −1900 and −1300 ms), the squared difference in mean pupil diameter during the preparatory phase predicted activity in the left middle frontal gyrus (MFG), extending into IFG (Fig. 4b; Supplementary Table 5). This relationship followed a U-shaped pattern (Fig. 4c). A similar right hemisphere source, in the IFG, showed a significant relationship with the squared difference in mean pupil diameter during the preparatory phase between −1400 and −1300 ms. These significant effects only occurred for squared pupil diameter, and no significant effects were found for signed pupil differences during the preparatory phase. As expected, we found no significant associations between preparatory-phase sources and either the raw or squared pupil differences measured during the selective phase.

To examine whether the significant association with squared pupil diameter was associated with any of the demographic or performance variables, we ran Bonferroni-corrected Pearson’s correlations examining activity in the MFG-IFG region of interest: Neither age, SRTs, accuracy, audiogram, nor squared transforms of these variables, correlated significantly with activity in this area (all |*r|* ≤ .32, all *p* ≥ .84); as expected, the signed difference in mean pupil diameter during the preparatory phase did not show a significant correlation (*r* = .30, *p* ∼ 1.00), whereas the squared difference in mean pupil diameter showed a strong positive correlation (*r* = .72, *p* < .001).

To check that the relationship with the squared difference in mean pupil diameter was not driven solely by the outlier shown in the top-right of Fig. 4e, we ran Pearson’s correlations excluding this participant. We found the same pattern of results with the remaining 29 participants: no significant correlation with the signed difference in mean pupil diameter during the preparatory phase (*r* = -.07, *p* = .71), and a significant positive correlation with the squared difference in pupil diameter (*r* = .43, *p* = .019).

When examining MEG sources during the selective phase, there were no significant effects.

## Discussion

Top-down attention is crucial for navigating complex environments, where multiple stimuli compete for perceptual processing. While robust group-level neural signatures of attentional control have been established in young adults, much less is known about how these vary across individuals, particularly in relation to sensory processing. Here, using a spatial cueing task, we combined MEG and pupillometry to characterise both shared and subject-specific aspects of top-down attention among older adults (Fig. 5). Across the group, we replicated established markers of attentional control, including sustained neural responses during preparation, alpha lateralisation, and recruitment of fronto-parietal and sensory regions. We also found that informative cues influenced target processing in auditory cortex and elicited greater pupil dilation than uninformative cues. Crucially, beyond these group-level effects, we observed systematic variability in cortical sources as a function of hearing acuity and pupil dilation—showing that attentional control is not uniform across individuals, but instead reflects individual differences in both sensory processing and strategic allocation of resources.

**Fig. 5.**
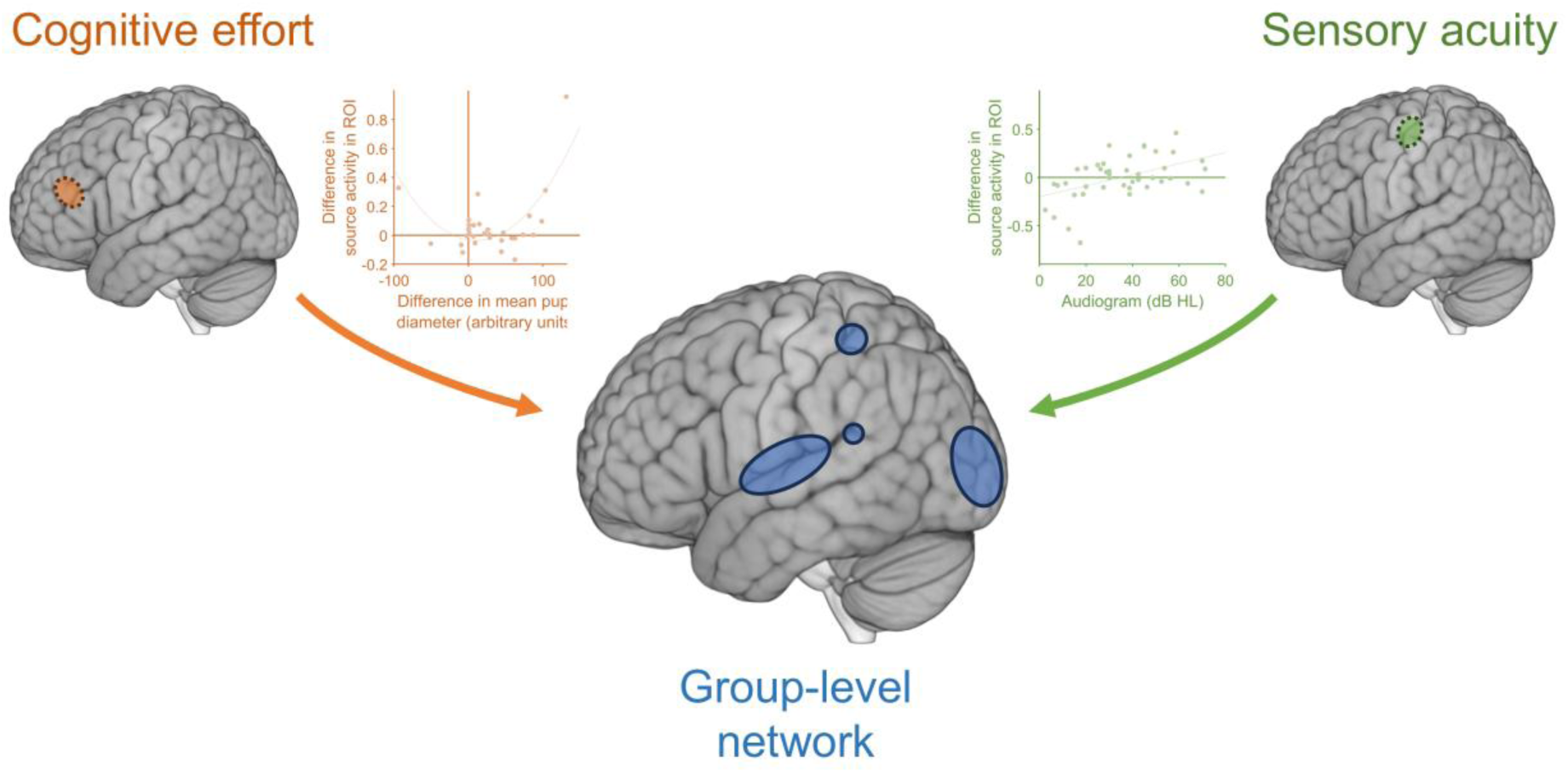
Schematic overview of main findings. Illustation of brain areas involved in preparatory attention. A core group-level network is supplemented by additional areas, which are differentially recruited depending on cognitive effort and sensory acuity. Here, cognitive effort and sensory acuity were found to have distinct—rather than overlapping—effects on the functional architecture for preparatory attention. Cognitive effort (indexed by pupil dilation) affected responses, non-linearly, in the left middle and inferior frontal gyri. Whereas lower sensory acuity (indexed by hearing thresholds) lead to greater responses in the left precentral and postcentral gyri.

These findings extend previous work in two key ways. First, by moving beyond group averages, they reveal that neural correlates of attentional control vary according to sensory processing—here, due to age-related hearing loss. Second, by linking neural activity to pupillometry, our results suggest that participants differ in their deployment of resources in preparation for a target stimulus—which is reflected in both differential pupil dilation and IFG engagement. Together, these results indicate that group-level responses only provide a partial account of top-down attention, and a fuller picture of the (degenerate) functional architecture is gained by incorporating these individual differences.

Interestingly, sensory loss was not associated with weaker engagement of attentional networks, but rather with *greater* responses to informative cues—specifically, in the left precentral and postcentral gyri. This result is consistent with compensatory recruitment of additional brain areas, which may increase cognitive demands, and could explain why individuals with greater sensory loss experience higher levels of fatigue^66^. Such recruitment is unlikely to fully restore performance to typical levels, because older adults who have more age-related hearing loss benefit less from spatial cues^15^. Notably, these effects did not localise to cingulo-opercular regions often implicated in ageing compensation^23–25^, but instead localised to precentral and postcentral gyri. One interpretation is that this activity reflects increased engagement of the frontal eye fields, which have been associated with successful spatial orienting^44,67,68^. While traditionally located more anteriorly^69^, the location of the frontal eye fields varies widely across studies, and human neuroimaging has implicated the precentral sulcus, extending into precentral gyrus^70^. Furthermore, a human MEG study^44^ identified the frontal eye fields by a functional localiser, and found that it extended from dorsal to ventral precentral gyrus in the left hemisphere—potentially overlapping with the area identified in the present study. Another possibility is that this effect reflects preparatory activation of motor areas involved in speech articulation, which are more strongly recruited under challenging listening conditions^71,72^. In addition, similar precentral and postcentral locations—along with the IFG—respond to visual cues that convey the content of upcoming speech^73^, and these areas may play a similar role during preparation in the current study. An alternative interpretation based on motor preparation for the button-press response is unlikely, because participants could not determine the correct response until after target onset. Overall, although precise localisation with MEG is limited, all of the most likely interpretations converge on the view that sensory loss shapes compensatory strategies in response to the spatial cues.

Compensatory strategies could depend on the age at which sensory loss occurs. Prior work demonstrated that children with early-onset hearing loss show weaker neural signatures of top-down attention in a similar task^45^. This contrasts with the present findings in older adults with age-related hearing loss, who show consistent evoked potentials to informative cues and greater recruitment of the precentral and postcentral gyri with increasing hearing loss. A likely explanation is that older adults—who experience gradual declines in hearing with older age—typically have a lifetime of experience with sounds and deploying spatial attention, such that they attempt to compensate for reduced sensory acuity. Taken together, these findings suggest that the consequences of sensory loss for attention depend on when in life the sensory loss occurred.

Pupil dilation is commonly interpreted as an index of arousal and cognitive effort and, here, it varied with the strength of IFG responses during the preparatory phase. One interpretation is that the IFG supports strategic allocation of resources, consistent with its involvement in the cingulo-opercular network^23–25^ and with previous interpretations of its role in effortful speech comprehension^27,29,74^. Such responses may correspond to the so-called ‘intensity’ of attention, which has been previously related to pupil responses and described as ‘mental effort’, separate from selective attention^42^. The relationship we found was non-linear, consistent with proposals that pupil diameter has a U-shaped relationship with performance^75–78^. The area that covaried with pupil dilation also included the MFG, which has previously been implicated in challenging speech perception in older adults^24^, and is related to pupil dilation while participants engage working memory^79^, which may be an additional component related to spatial cueing in the present task.

Our results underscore that the functional architecture evoked by spatial cues is shaped by multiple factors, which opens new potential avenues for investigation. For example, future work could examine how preparatory processes interact with other factors, such as working memory and domain-general cognitive processes. More broadly, our results imply that a group-level approach may be suboptimal for interventions aimed at enhancing attention, as individuals vary in the neural strategies they deploy for the same task. Instead, interventions may be most effective when tailored to an individual’s sensory profile and attentional strategy. While current research efforts have considered personalised interventions for hearing^80^, integrating information about an individual’s functional architecture may improve their efficacy—a possibility that warrants systematic investigation.

In summary, our findings demonstrate that top-down attention is not a uniform process but instead reflects systematic individual differences shaped by sensory processing and strategic resource allocation. By combining MEG and pupillometry, we show that group-level signatures of attention capture only part of the story, and that considering heterogenous, degenerate functional architectures provides a richer account of attentional control.

## Methods

### Participants

We analysed data from 44 participants. We conducted power analyses using G*Power^81^ (version 3.1.9.7), which showed that a sample size of 44 is sensitive to correlations (e.g., between MEG activity and audiometric thresholds) of size *r^2^*≥ .16 (alpha = .05, power = .8). An additional 7 participants were recruited but did not complete the experiment: 5 participants had strong dental artifacts in their MEG data at rest, so did not continue; one participant chose to withdraw; another did not complete the experiment in the available time.

The 44 participants (25 female, 19 male; 36 right-handed, 5 left-handed, 3 ambidextrous) were aged 56–80 years (median = 66.5 years, inter-quartile range = 9.3). They were all native English speakers with normal or corrected-to-normal vision. None of the participants used cochlear implants or had any history of ear surgery. Two participants reported experiencing tinnitus. Participants who used hearing aids (N = 2) used their hearing aids while listening to the instructions, but removed them before starting the experimental tasks.

We measured participants’ pure-tone audiometric thresholds at octave frequencies between 0.25 and 8 kHz using a Starkey Acoustic Analyser AA30 (Starkey Laboratories, Inc.) in accordance with BS EN ISO 8253-124^82^. Participants had pure-tone average thresholds, averaged across ears, of .8–47.9 dB HL (median = 17.9 dB HL, inter-quartile range = 12.7), with a difference of .0–11.7 dB HL (median = 4.2 dB HL, inter-quartile range = 3.3) between the left and right ears. The pure-tone audiometric thresholds from individual participants are displayed in Fig. 3c.

The study was approved by the [ethics committee and ID number], and was performed in accordance with relevant guidelines and regulations. Informed consent was obtained from all participants.

### Equipment

The experiment was conducted in a magnetically shielded room containing a CTF-275 MEG system (axial gradiometers; 274 channels; 30 reference channels; VSM MedTech Ltd., Coquitlam, Canada) and an Eyelink 1000 eye-tracker (SR Research Ltd., Ottawa, Canada). The lights in the room were turned off during recording.

Visual stimuli were presented on an JVC DLA-SX21 (JVC Professional Products Company, New Jersey, USA) projector, placed at a distance of approximately 55 cm in front of participants’ eyes. Acoustic stimuli were delivered using MEG-compatible earphones (Etymotic Ear-Tone ER3A, Illinois, USA). Participants responded using a button box positioned under their right hand.

Practice trials were completed in a sound-attenuating booth outside of the scanner area. Participants sat in a comfortable chair facing an LCD visual display unit (Dell Technologies, Texas, USA). Acoustic stimuli were presented through an external sound card (ESI Maya 22 USB; ESI Audiotechnik GmbH, Leonberg) connected to circumaural headphones (Sennheiser HD 380 Pro; Sennheiser electronic GmbH & Co. KG, Wedemark, Germany). Participants responded using 4 buttons on a computer keyboard.

All tasks were delivered with Psychtoolbox^81^ (version: 3.0.17) using custom-written MATLAB scripts (version R2019a; MathWorks, Inc., Natick, USA).

### Stimuli

Acoustic stimuli were modified phrases from the Co-ordinate Response Measure corpus^83^ (CRM), used by Holmes et al.^45,49^. Each phrase contained a colour word (“Red”, “Green”, “Blue” or “White”) and a number word (“one”, “two”, “three” or “four”), followed by the word “now”. For example, “Red two now”. The phrases were spoken by native British English talkers. We used one male talker, one female talker, and a “child” talker, which was the voice of a different female talker that was manipulated using Praat (Version 5.3.08; http://www.praat.org/) to have a higher fundamental frequency and shorter formant spacing ratio than the original recordings. The digital recordings had an average duration of 1.4 seconds and were adjusted to have equal root mean square (RMS) amplitudes.

Visual cues were white arrows that pointed leftwards or rightwards on a grey background. A visual composite stimulus, which was used as the uninformative cue, contained both arrows overlaid, on a grey background. We also used a fixation cross stimulus, which was a white cross on a grey background. The images were adjusted to have equal intensities. Afterwards, a white rectangle was added to the lower left corner of each image; this rectangle was not visible to participants (as it was projected off-screen), but allowed an accurate measurement of the stimulus presentation times through a photodiode signal recorded as a channel in the MEG acquisition.

### Procedure

Fig. 1 illustrates the trial structure for the spatial attention task. On every trial, participants saw a fixation cross, then a pre-cue, followed by a fixation cross, and then a left or right spatial cue, which indicated the location of the target talker and varied from trial to trial. There were two pre-cue conditions: In the *uninformative* pre-cue condition, the pre-cue was the composite stimulus that did not inform the participant about the target location; in the *informative* pre-cue condition, the pre-cue was a preview of the spatial cue (that was always valid).

The acoustic stimuli for each trial were three CRM phrases overlaid with equal RMS amplitudes. The three phrases presented on each trial were always spoken by different talkers, contained different colour and number words, and were simulated to come from different spatial locations using interaural time differences (ITDs). The child’s voice was always presented with an ITD of 0 µs; in other words, it was simulated to originate from a central location and was never the target. The locations of the male and female voices varied pseudo-randomly on each trial. One had an ITD of −205 µs (i.e., simulated to the left side), and the other had an ITD of +205 µs, (i.e., simulated to the right side). After the acoustic stimuli had ended (and 3.2 seconds after their onset), participants were first required to report the colour word (“Red”, “Green”, “Blue” or “White”) from the target sentence by pressing one of four buttons. Then, they were asked to report the number word they heard (“one”, “two”, “three” or “four”) by pressing one of the same four buttons. The intertrial interval was randomly selected on each trial, between 1.5 and 2.5 seconds.

Before entering the scanner area, participants completed four practice trials of the spatial attention task. Practice trials had the same format as the main task, except that explicit feedback (“Correct” or “Incorrect”) was provided after each trial. Participants were allowed to repeat the practice trials as many times as they wanted, until they felt that they had a good understanding of the task.

Once participants were positioned in the MEG scanner, they first completed a speech reception threshold (SRT) task, in which the overall level of individual CRM phrases were adapted to each participant’s 50% threshold in quiet. On each trial, participants first saw a fixation cross, which was displayed for 1000 ms before the speech began. They then heard a single CRM phrase, which could be spoken by any of the three talkers and was presented with an ITD of 0 µs. Participants were prompted to make a response 500 ms after the phrase had ended. Like in the spatial attention task, participants were first asked to report the colour word from the CRM phrase, by pressing one of four buttons, and then the number word, by pressing one of the same four buttons. Participants were instructed to keep their eyes open during each trial and fixate at the centre of the fixation cross. We varied the level of the acoustic stimuli in a 1-up 1-down adaptive procedure^84^. Each run started at a level of 70 dB(A) SPL. The step size started at 5 dB and decreased to 1 dB after 3 reversals. For each task, we adapted the level in two separate but interleaved runs; the two runs were structured in the same way, except different stimuli were presented. Each run terminated after 10 reversals. The SRT for each run was calculated as the median of the last 6 reversals, and the SRT for each participant was calculated as the average of the SRTs across the two runs.

Participants then completed 240 trials of the spatial attention task, which were separated into 10 blocks of 24 trials. Each block contained 6 trials from each condition (2 pre-cue conditions x 2 spatial cues). The combined level of the three CRM phrases presented on each trial were set to 25 dB above the participant’s SRT, which participants reported to be a comfortable listening level. No feedback was given after each trial. Participants were instructed to keep their eyes open during each trial and fixate at the centre of the screen.

Finally, 33 participants completed a passive visual control task, in which the visual stimuli were presented, but there were no acoustic stimuli and participants did not make any responses. Seven participants did not complete this part due to time constraints. On each trial of the visual orienting block, participants saw a fixation cross for 1000 ms, followed by either the leftwards-pointing arrow, the rightwards-pointing arrow, or the visual composite stimulus, which lasted for 300 ms. The intertrial interval was randomly selected on each trial, between 3.5 and 4.5 seconds. Participants completed 45 trials (15 for each of the three visual stimuli). They were instructed to keep their eyes open during each trial and fixate at the centre of the screen.

### MEG Recording and Pre-Processing

MEG data were recorded at a sampling rate of 1200 Hz. We conducted all analyses in MATLAB (version R2024b; MathWorks, Inc., Natick, USA). We pre-processed the MEG data (from gradiometer sensors) using the SPM12 toolbox (Wellcome Centre for Human Neuroimaging, London, UK).

For analysing magnetic evoked fields and conducting source reconstruction for the spatial attention task, we epoched the data at −2500 to 1800 ms, relative to the onset of acoustic stimuli. The timing of each epoch was adjusted to account for a 107.5 ms delay between the triggers and stimulus presentation (as indicated by the photodiode recording). We applied subtractive baseline correction, using a baseline of −2500 to −2300 ms (i.e., 200 ms at the end of the fixation cross period, before the pre-cue was presented). Given we were interested in individual differences in brain responses, we excluded incorrect trials from the analyses, so that differences across participants reflect differences on successful trials, without confounding differences in accuracy across participants. (However, we also analysed magnetic evoked fields on incorrect trials, and found no significant differences between correct and incorrect trials; cluster-based permutation analysis: sum of *t*-values ≤ 350.56, *p* ≥ 0.19). We also rejected trials containing high-amplitude artifacts (using a peak-to-peak amplitude threshold of 3000). We performed robust averaging across trials, then applied a low-pass filter at 60 Hz to correct for any high-frequency noise introduced by the averaging procedure. Finally, we calculated grand means across participants.

For plotting magnetic evoked fields, we calculated the root-mean-square (RMS) amplitude across all channels. Given evoked fields were similar for left and right cued conditions (Supplementary Fig. 1a), we collapsed across left and right cue types to compare informative and uninformative pre-cue conditions.

To prepare the data for time-frequency analyses, we followed a similar pre-processing pipeline, except that we defined a longer duration for epochs (−2800 to 2100 ms). We applied a multi-taper spectral decomposition at 6–40 Hz, and used a logistic baseline correction, such that the time-frequency representation reflects decibel changes from baseline. To isolate alpha activity, we averaged across 8–12 Hz, based on previous studies^85^.

When analysing data from the passive visual control task, we used an epoch window of −500 to 2000 ms, relative to the onset of visual stimuli. We used a baseline of −500 to −300 ms (i.e., 200 ms during fixation cross presentation). Given participants did not make any responses during the passive visual control task, there was no performance-based exclusion of trials.

### MEG Source Localisation

We used SPM12 (Wellcome Centre for Human Neuroimaging, London, UK) for source localisation, which we conducted on the magnetic evoked fields. We co-registered the data to MNI coordinates based on the nasion, and left and right preauricular fiducials. We then conducted a group inversion (using the data from all participants) using a single shell forward model and multiple sparse priors (greedy search method) for source inversion. We conducted the source analysis across the entire epoch window, then inspected the results within our time windows of interest.

For the spatial attention task, we analysed responses during the preparatory and selective phases. For the preparatory phase, our time window was −2000 to −300 ms, which corresponded to the period of time that the fixation cross was on the screen, so the visual stimuli were identical in all conditions, but participants had knowledge of the upcoming cue in the informative condition and not in the uninformative condition. For the selective phase, we used a time window of 0 to 1000 ms. These time windows should be sensitive to low-frequency responses and sources that display sustained activity throughout each phase. To examine time-varying responses in the preparatory phase, we also inspected results within 100-ms non-overlapping sub-windows (e.g., −2000 to −1900 ms, −1900 to −1800 ms, etc.).

For the passive visual control task, we analysed responses at 0 to 1700 ms (relative to the visual stimulus onset), to match the duration of time to the 1700-ms window used for the preparatory phase of the spatial attention task.

### Eye-Tracker Recording and Pre-Processing

We analysed simultaneous eye-tracking data from 30 participants. For the remaining 14 participants, participants’ pupils were not visible from the eye-tracking camera (because the position of the MEG helmet obscured the view; N = 9) or the pupil data were excessively noisy (N = 5). The 30 remaining participants (13 male, 17 female; 23 right-handed, 4 left-handed, 3 ambidextrous) were aged 56–79 years (median = 65.5 years; interquartile range = 8.8).

The eye-tracker was calibrated for each participant using 9-point calibration. Eye-tracking data were recorded from one eye at a sample rate of either 500 Hz (N = 11) or 1000 Hz (N = 19). We imported the EyeLink files using the Edf2Mat MATLAB Toolbox (designed and developed by Adrian Etter and Marc Biedermann at the University of Zurich), then pre-processed the data using the Pupillometry Pipeliner toolbox^86^. We identified blinks using the pupillometry noise method^87^, then removed data surrounding the blink (−50 to 150 ms) and interpolated the removed data (using linear interpolation, with a maximum duration of 600 ms). We smoothed the data using a median moving window (with a width of 150 ms). We then epoched the data between −3300 ms and 5700 ms and subtracted the mean from a baseline window of −2800 to −2300 ms (i.e., 500 ms at the end of the fixation cross period, before the pre-cue was presented). We rejected epochs in which > 20% of the data were missing. Where the sample rate was greater than 500 Hz (N = 19), we down-sampled the data, so that all participants’ data were sampled at 500 Hz.

To verify that differences in pupil diameter between conditions could not be explained by eye movements—which can result in smaller apparent pupil diameter due to the foreshortening error^88,89^—we re-analysed the data after applying the pupil foreshortening error correction^90^ and we observed the same pattern of results (Supplementary Fig. 3) as in the main analysis (Fig. 4a).

### Statistical Analyses

To analyse behavioural data for the spatial attention task, we performed a two-way repeated measures ANOVA (informative/uninformative pre-cues and attend-left/attend-right trials).

To analyse differences in magnetic evoked fields between informative and uninformative pre-cue conditions, we conducted a cluster-based permutation analysis on the RMS amplitude across all channels. This analysis was conducted in the time-domain, across the entire epoch window. We performed a two-sided test for dependent samples using the ‘permutest’ function^91^ for MATLAB. We set the *p*-threshold (for determining which data points belonged to the same clusters) to .05 and derived *p*-values at the cluster-level from 10,000 permutations of the data. We repeated this analysis for the passive visual control task, comparing trials in which the left or right arrow stimuli (which corresponded to the informative pre-cues in the selective attention task) were presented to trials in which the visual composite stimulus (which corresponded to the uninformative pre-cue in the selective attention task) was presented.

To examine alpha lateralisation, we subtracted attend-right from attend-left trials and inspected the scalp maps; we included informative and uninformative trials when creating scalp maps for the selective phase, but we only included informative trials when creating scalp maps for the preparatory phase, given that there were no differences between attend-left and attend-right trials until after the preparatory phase had ended.

For source analyses, we subtracted the uninformative condition from the informative condition (collapsing across left and right cueing conditions) for each participant. Using a general linear model, we conducted one-sample *t*-tests in SPM12 (on each time window of interest), with four between-subjects covariates: age; speech reception thresholds (SRTs), accuracy in spatial attention task (collapsed across the four conditions), and audiometric thresholds (at 4–8 kHz, averaged across ears, given these frequencies are most sensitive to age-related hearing loss^11^).

To analyse pupil time-courses, we conducted a cluster-based permutation analysis (*p*-threshold = .05; 10,000 permutations) that assessed differences between informative and uninformative conditions across the group of participants. To link pupil diameter to MEG sources, we conducted additional statistical analyses on the MEG source images for the 30 participants with usable pupil data. Using a general linear model, we ran one-sample *t*-tests on the MEG source images (on each time window of interest) using four between-subjects pupil covariates. The covariates included the difference in mean pupil diameter between the informative and uninformative conditions in two time windows: one intended to capture differences during the preparatory phase (−2000 to 0 ms), and the second intended to capture differences during the selective phase (0 to 4000 ms). The remaining two covariates were squared differences in the two time windows, to capture non-linear effects^64,65^. We used longer time windows for the pupil analyses than the MEG analyses to account for delays between stimulus presentation and pupil dilation^40,92^.

Outliers were defined using three standard deviations from the mean. All time windows are relative to the onset of the acoustic stimuli.

## Data availability

The anonymised data generated during this study are available at the Open Science Framework ([OSF URL]).

## Code availability

The code used to run the experiment is available at the Open Science Framework ([OSF URL]). Data were analysed using standard MATLAB code available in the open-source SPM12 academic software (http://www.fil.ion.ucl.ac.uk/spm).

## Ethics declarations

The authors declare no competing interests.

## Acknowledgements

This study was funded by an RNID Pauline Ashley Fellowship (PA25_Holmes) to EH. The study was conducted at the Wellcome Centre for Human Neuroimaging, which was supported by Wellcome (091593/Z/10/Z). We thank Andrea Caso for assisting with participant recruitment and data collection.

## Notes

https://osf.io/3q7u6/overview

### Competing Interest Statement

The authors have declared no competing interest.

## References

1. Nobre, A. C. & Mesulam, M. M. Large-scale Networks for Attentional Biases. in The Oxford Handbook of Attention (eds. Nobre, A. C. & Kastner, S.) 105–151 (Oxford University Press, 2014). doi:10.1093/OXFORDHB/9780199675111.001.0001.

2. Hill, K. T. & Miller, L. M. Auditory attentional control and selection during cocktail party listening. Cereb. Cortex 20, 583–90 (2010).

3. Woldorff, M. G. et al. Functional parcellation of attentional control regions of the brain. J. Cogn. Neurosci. 16, 149–65 (2004).

4. Hu, M. et al. Neural processes of auditory perception in Heschl’s gyrus for upcoming acoustic stimuli in humans. Hear. Res. 388, (2020).

5. Sajid, N., Parr, T., Hope, T. M., Price, C. J. & Friston, K. J. Degeneracy and Redundancy in Active Inference. Cereb. Cortex 30, 5750–5766 (2019).

6. Price, C. J. & Friston, K. J. Degeneracy and cognitive anatomy. Trends Cogn. Sci. 6, 416–421 (2002).

7. Tononi, G., Sporns, O. & Edelman, G. M. Measures of degeneracy and redundancy in biological networks. Proc. Natl. Acad. Sci. U. S. A. 96, 3257–3262 (1999).

8. Holmes, E. & Griffiths, T. D. ‘Normal’ hearing thresholds and fundamental auditory grouping processes predict difficulties with speech-in-noise perception. Sci. Rep. 9, 16771 (2019).

9. Motlagh Zadeh, L., et al. Extended high-frequency hearing enhances speech perception in noise. Proc. Natl. Acad. Sci. U. S. A. 116, 23753–23759 (2019).

10. Brant, L. J. & Fozard, J. L. Age changes in pure-tone hearing thresholds in a longitudinal study of normal human aging. J. Acoust. Soc. Am. 88, 813–820 (1990).

11. Wiley, T. L., Chappell, R., Carmichael, L., Nondahl, D. M. & Cruickshanks, K. J. Changes in hearing thresholds over 10 years in older adults. J Am Acad Audiol. 19, 281–371 (2008).

12. Killeen, O. J., Zhou, Y. & Ehrlich, J. R. Objectively measured visual impairment and dementia prevalence in older adults in the US. JAMA Ophthalmol. 141, 786–790 (2023).

13. Loughrey, D. G., Kelly, M. E., Kelley, G. A., Brennan, S. & Lawlor, B. A. Association of age-related hearing loss with cognitive function, cognitive impairment, and dementia a systematic review and meta-analysis. JAMA Otolaryngol. - Head Neck Surg. 144, 115–126 (2018).

14. Livingston, G. et al. Dementia prevention, intervention, and care: 2020 report of the Lancet Commission. Lancet 396, 413–446 (2020).

15. Caso, A., Griffiths, T. D. & Holmes, E. Spatial selective auditory attention is preserved in older age but is degraded by peripheral hearing loss. Sci. Rep. 14, 26243 (2024).

16. Macsweeney, M., Brammer, M. J., Waters, D. & Goswami, U. Enhanced activation of the left inferior frontal gyrus in deaf and dyslexic adults during rhyming. Brain 132, 1928–1940 (2009).

17. Baker, C. I., Peli, E., Knouf, N. & Kanwisher, N. G. Reorganization of visual processing in macular degeneration. J. Neurosci. 25, 614–618 (2005).

18. Kanai, R., Komura, Y., Shipp, S. & Friston, K. J. Cerebral hierarchies: predictive processing, precision and the pulvinar. Philos. Trans. R. Soc. B Biol. Sci. 370, 20140169–20140169 (2015).

19. Oosterhuis, E. J., Slade, K., May, P. J. C. & Nuttall, H. E. Toward an Understanding of Healthy Cognitive Aging: The Importance of Lifestyle in Cognitive Reserve and the Scaffolding Theory of Aging and Cognition. Journals Gerontol. Ser. B 78, 777–788 (2023).

20. Phillips, N. A. The Implications of Cognitive Aging for Listening and the Framework for Understanding Effortful Listening (FUEL). Ear Hear. 37, 44S–51S (2016).

21. Dahl, M. J., Ilg, L., Li, S. C., Passow, S. & Werkle-Bergner, M. Diminished pre-stimulus alpha-lateralization suggests compromised self-initiated attentional control of auditory processing in old age. Neuroimage 197, 414–424 (2019).

22. Hong, X., Sun, J., Bengson, J. J., Mangun, G. R. & Tong, S. Normal aging selectively diminishes alpha lateralization in visual spatial attention. Neuroimage 106, 353–363 (2015).

23. Eckert, M. A. et al. Age-related effects on word recognition: Reliance on cognitive control systems with structural declines in speech-responsive cortex. JARO - J. Assoc. Res. Otolaryngol. 9, 252–259 (2008).

24. Erb, J. & Obleser, J. Upregulation of cognitive control networks in older adults’ speech comprehension. Front. Syst. Neurosci. 7, 116 (2013).

25. Harris, K. C., Dubno, J. R., Keren, N. I., Ahlstrom, J. B. & Eckert, M. A. Speech recognition in younger and older adults: a dependency on low-level auditory cortex. 29, 6078–6087 (2009).

26. Wingfield, A. & Grossman, M. Language and the aging brain: Patterns of neural compensation revealed by functional brain imaging. J. Neurophysiol. 96, 2830–2839 (2006).

27. Vaden Jr, K. I., Kuchinsky, S. E., Ahlstrom, J. B., Dubno, J. R. & Eckert, M. a. Cortical Activity Predicts Which Older Adults Recognize Speech in Noise and When. J. Neurosci. 35, 3929–3937 (2015).

28. Peelle, J. E., Troiani, V., Wingfield, A. & Grossman, M. Neural processing during older adults’ comprehension of spoken sentences: Age differences in resource allocation and connectivity. Cereb. Cortex 20, 773–782 (2010).

29. Vaden Jr, K. I., et al. The cingulo-opercular network provides word-recognition benefit. J. Neurosci. 33, 18979–18986 (2013).

30. Kerns, J. G. Anterior cingulate and prefrontal cortex activity in an FMRI study of trial-to-trial adjustments on the Simon task. Neuroimage 33, 399–405 (2006).

31. Kerns, J. G. et al. Anterior cingulate conflict monitoring and adjustments in control. Science (80-. ). 303, 1023–1027 (2004).

32. Walsh, B. J., Buonocore, M. H., Carter, C. S. & Mangun, G. R. Integrating Conflict Detection and Attentional Control Mechanisms. J. Cogn. Neurosci. 23, 2211 (2011).

33. Alavash, M. & Obleser, J. Brain network interconnectivity dynamics explain metacognitive differences in listening behavior. J. Neurosci. 44, e2322232024 (2024).

34. Botvinick, M. M., Cohen, J. D. & Carter, C. S. Conflict monitoring and anterior cingulate cortex: an update. Trends Cogn. Sci. 8, 539–46 (2004).

35. Winn, M. B., Edwards, J. R. & Litovsky, R. Y. The impact of auditory spectral resolution on listening effort revealed by pupil dilation. Ear Hear. 36, e135–e165 (2015).

36. Zekveld, A. A., Kramer, S. E. & Festen, J. M. Cognitive load during speech perception in noise: The influence of age, hearing loss, and cognition on the pupil response. Ear Hear. 32, 498–510 (2011).

37. Murphy, P. R., O’Connell, R. G., O’Sullivan, M., Robertson, I. H. & Balsters, J. H. Pupil diameter covaries with BOLD activity in human locus coeruleus. Hum. Brain Mapp. 35, 4140–4154 (2014).

38. de Gee, J. W. et al. Dynamic modulation of decision biases by brainstem arousal systems. Elife 6, (2017).

39. Meissner, S. N. et al. Self-regulating arousal via pupil-based biofeedback. Nat. Hum. Behav. 2023 81 8, 43–62 (2023).

40. Reimer, J. et al. Pupil fluctuations track rapid changes in adrenergic and cholinergic activity in cortex. Nat. Commun. 7, 1–7 (2016).

41. Jones, B. E. Activity, modulation and role of basal forebrain cholinergic neurons innervating the cerebral cortex. Prog. Brain Res. 145, 157–169 (2004).

42. Unsworth, N. & Miller, A. L. Individual Differences in the Intensity and Consistency of Attention. Curr. Dir. Psychol. Sci. 30, 391–400 (2021).

43. Piquado, T., Isaacowitz, D. & Wingfield, A. Pupillometry as a measure of cognitive effort in younger and older adults. Psychophysiology 47, 560–569 (2010).

44. Lee, A. K. C. et al. Auditory selective attention reveals preparatory activity in different cortical regions for selection based on source location and source pitch. Front. Neurosci. 6, 1–9 (2013).

45. Holmes, E., Kitterick, P. T. & Summerfield, A. Q. Peripheral hearing loss reduces the ability of children to direct selective attention during multi-talker listening. Hear. Res. 350, 160–172 (2017).

46. Cherry, E. C. Some experiments on the recognition of speech, with one and two ears. J. Acoust. Soc. Am. 25, 1262–2527 (1953).

47. Corbetta, M., Kincade, J. M., Ollinger, J. M., McAvoy, M. P. & Shulman, G. L. Voluntary orienting is dissociated from target detection in human posterior parietal cortex. Nat. Neurosci. 3, 292–7 (2000).

48. Hopfinger, J. B., Buonocore, M. H. & Mangun, G. R. The neural mechanisms of top-down attentional control. Nat. Neurosci. 3, 284–91 (2000).

49. Holmes, E., Kitterick, P. T. & Summerfield, A. Q. Cueing listeners to attend to a target talker progressively improves word report as the duration of the cue-target interval lengthens to 2,000 ms. *Attention, Perception*, Psychophys. 80, 1520–1538 (2018).

50. Alavash, M., Tune, S. & Obleser, J. Dynamic Large-Scale Connectivity of Intrinsic Cortical Oscillations Supports Adaptive Listening in Challenging Conditions. PLoS Biology vol. 19 (2021).

51. Tune, S., Alavash, M., Fiedler, L. & Obleser, J. Neural attentional-filter mechanisms of listening success in middle-aged and older individuals. Nat. Commun. 12, 1–14 (2021).

52. Best, V., Marrone, N., Mason, C. R., Kidd, G. & Shinn-Cunningham, B. G. Effects of sensorineural hearing loss on visually guided attention in a multitalker environment. J. Assoc. Res. Otolaryngol. 10, 142–9 (2009).

53. Dai, L., Best, V. & Shinn-Cunningham, B. G. Sensorineural hearing loss degrades behavioral and physiological measures of human spatial selective auditory attention. Proc. Natl. Acad. Sci. 115, E3286–E3295 (2018).

54. Holmes, E., Kitterick, P. T. & Summerfield, A. Q. EEG activity evoked in preparation for multi-talker listening by adults and children. Hear. Res. 336, 83–100 (2016).

55. Wöstmann, M., Herrmann, B., Maess, B. & Obleser, J. Spatiotemporal dynamics of auditory attention synchronize with speech. Proc. Natl. Acad. Sci. 113, 3873–3878 (2016).

56. Kerlin, J. R., Shahin, A. J. & Miller, L. M. Attentional gain control of ongoing cortical speech representations in a “cocktail party”. J. Neurosci. 30, 620–628 (2010).

57. ElShafei, H. A., Bouet, R., Bertrand, O. & Bidet-Caulet, A. Two Sides of the Same Coin: Distinct Sub-Bands in the Alpha Rhythm Reflect Facilitation and Suppression Mechanisms during Auditory Anticipatory Attention. Eneuro ENEURO.0141-18.2018 (2018) doi:10.1523/ENEURO.0141-18.2018.

58. Humes, L. E., Kidd, G. R. & Lentz, J. J. Auditory and cognitive factors underlying individual differences in aided speech-understanding among older adults. Front Syst Neurosci 7, 55 (2013).

59. van Rooij, J. C. G. M. & Plomp, R. Auditive and cognitive factors in speech perception by elderly listeners. III. Additional data and final discussion. J. Acoust. Soc. Am. 91, 1028–1033 (1992).

60. Helfer, K. S. & Freyman, R. L. Stimulus and listener factors affecting age-related changes in competing speech perception. J. Acoust. Soc. Am. 136, 748 (2014).

61. Goossens, T., Vercammen, C., Wouters, J. & van Wieringen, A. Masked speech perception across the adult lifespan: Impact of age and hearing impairment. Hear. Res. 344, 109–124 (2017).

62. Scimeca, J. M., Kiyonaga, A. & D’Esposito, M. Reaffirming the sensory recruitment account of working memory. Trends in Cognitive Sciences vol. 22 190–192 at 10.1016/j.tics.2017.12.007 (2018).

63. Schwalm, M. & Rosales Jubal, E. Back to pupillometry: How cortical network state fluctuations tracked by pupil dynamics could explain neural signal variability in human cognitive neuroscience. eNeuro 4, 1–9 (2017).

64. Kaufman, K. J., Krall, R. F. & Williamson, R. S. Pupil-linked arousal differentially modulates cell-type-specific sensory processing. *bioRxiv* 2025.06.09.658645 (2025) doi:10.1101/2025.06.09.658645.

65. Schwartz, Z. P., Buran, B. N. & David, S. V. Pupil-associated states modulate excitability but not stimulus selectivity in primary auditory cortex. J. Neurophysiol. 123, 191–208 (2020).

66. Jiang, K., Spira, A. P., Lin, F. R., Deal, J. A. & Reed, N. S. Hearing Loss and Fatigue in Middle-Aged and Older Adults. JAMA Otolaryngol. - Head Neck Surg. 149, 758–760 (2023).

67. Bollimunta, A., Bogadhi, A. R. & Krauzlis, R. J. Comparing frontal eye field and superior colliculus contributions to covert spatial attention. Nat. Commun. 9, 3553 (2018).

68. Moore, T. & Armstrong, K. M. Selective gating of visual signals by microstimulation of frontal cortex. Nat. 2003 4216921 421, 370–373 (2003).

69. Tehovnik, E. J., Sommer, M. A., Chou, I. H., Slocum, W. M. & Schiller, P. H. Eye fields in the frontal lobes of primates. Brain Res. Rev. 32, 413–448 (2000).

70. Vernet, M., Quentin, R., Chanes, L., Mitsumasu, A. & Valero-Cabré, A. Frontal eye field, where art thou? Anatomy, function, and non-invasive manipulation of frontal regions involved in eye movements and associated cognitive operations. Front. Integr. Neurosci. 8, 66 (2014).

71. Cope, T. E. et al. Temporal lobe perceptual predictions for speech are instantiated in motor cortex and reconciled by inferior frontal cortex. Cell Rep. 42, 112422 (2023).

72. Perron, M., Vuong, V., Grassi, M. W., Imran, A. & Alain, C. Engagement of the speech motor system in challenging speech perception: Activation likelihood estimation meta-analyses. Hum. Brain Mapp. 45, e70023 (2024).

73. Sohoglu, E., Peelle, J. E., Carlyon, R. P. & Davis, M. H. Predictive top-down integration of prior knowledge during speech perception. J. Neurosci. 32, 8443–53 (2012).

74. Wild, C. J. et al. Effortful listening: the processing of degraded speech depends critically on attention. J. Neurosci. 32, 14010–21 (2012).

75. Murphy, P. R., Robertson, I. H., Balsters, J. H. & O’connell, R. G. Pupillometry and P3 index the locus coeruleus-noradrenergic arousal function in humans. Psychophysiology 48, 1532–1543 (2011).

76. Wendt, D., Koelewijn, T., Książek, P., Kramer, S. E. & Lunner, T. Toward a more comprehensive understanding of the impact of masker type and signal-to-noise ratio on the pupillary response while performing a speech-in-noise test. Hear. Res. 369, 67–78 (2018).

77. Ohlenforst, B. et al. Impact of SNR, masker type and noise reduction processing on sentence recognition performance and listening effort as indicated by the pupil dilation response. Hear. Res. 365, 90–99 (2018).

78. Hauswald, A., Keitel, A., Chen, Y.-P. Y., Rösch, S. & Weisz, N. Degradation levels of continuous speech affect neural speech tracking and alpha power differently. Eur. J. Neurosci. ejn.14912 (2020).

79. Siegle, G. J., Steinhauer, S. R., Stenger, V. A., Konecky, R. & Carter, C. S. Use of concurrent pupil dilation assessment to inform interpretation and analysis of fMRI data. Neuroimage 20, 114–124 (2003).

80. Sanchez-Lopez, R., Fereczkowski, M., Santurette, S., Dau, T. & Neher, T. Towards Auditory Profile-Based Hearing-Aid Fitting: Fitting Rationale and Pilot Evaluation. Audiol. Res. 11, 10 (2021).

81. Faul, F., Erdfelder, E., Lang, A.-G. & Buchner, A. G*Power 3: A flexible statistical power analysis program for the social, behavioral, and biomedical sciences. Behav. Res. Methods 39, 175–191 (2007).

82. British Society of Audiology. Recommended Procedure: Pure Tone Air and Bone Conduction Threshold Audiometry with and without Masking and Determination of Uncomfortable Loudness Levels. http://www.thebsa.org.uk/docs/bsapta.doc (2004).

83. Moore, T. J. Voice communication jamming research. in AGARD Conference Proceedings 331: Aural Communication in Aviation 2:1-2:6 (Neuilly-Sur-Seine, France, 1981).

84. Cornsweet, T. N. The Staircase-Method in psychophysics. Am. J. Psychol. 75, 485–491 (1962).

85. Wöstmann, M., Alavash, M. & Obleser, J. Alpha oscillations in the human brain implement distractor suppression independent of target selection. J. Neurosci. 1954–19 (2019) doi:10.1523/jneurosci.1954-19.2019.

86. Kinley, I. & Levy, Y. PuPl: an open-source tool for processing pupillometry data. Behav. Res. Methods 54, 2046–2069 (2022).

87. Hershman, R., Henik, A. & Cohen, N. CHAP: Open-source software for processing and analyzing pupillometry data. Behav. Res. Methods 51, 1059–1074 (2019).

88. Hayes, T. R. & Petrov, A. A. Mapping and correcting the influence of gaze position on pupil size measurements. Behav. Res. Methods 48, 510–527 (2016).

89. Gagl, B., Hawelka, S. & Hutzler, F. Systematic influence of gaze position on pupil size measurement: Analysis and correction. Behav. Res. Methods 43, 1171–1181 (2011).

90. Brisson, J. et al. Pupil diameter measurement errors as a function of gaze direction in corneal reflection eyetrackers. Behav. Res. Methods 45, 1322–1331 (2013).

91. Gerber, E. M. permutest. at (2024).

92. Wierda, S. M., Van Rijn, H., Taatgen, N. A. & Martens, S. Pupil dilation deconvolution reveals the dynamics of attention at high temporal resolution. Proc. Natl. Acad. Sci. U.S. A. 109, 8456–8460 (2012).

