## Supplementary for "Heterogeneity in neural network engagement supports individual differences in top-down attention"

**Supplementary Fig. 1.** Group-level results. **a**, Magnetic evoked fields across all participants (N = 44) in the informative and uninformative conditions, separated into attend-left and attend-right trials. Lines show the means and standard errors. **b**, Magnetic evoked fields in the passive visual control task, across all participants who completed this task (N = 33). Lines show the means and standard errors for trials in which the informative cue stimuli (left and right arrows) and uninformative cue stimulus (composite stimulus) were displayed on the screen. **c**, Sensor maps showing alpha activity (8–12 Hz) for the difference between attend-left and attend-right trials during the preparatory and selective phases (N = 44). **d**, Source analysis during the selective phase (0 to 1000 ms), showing the difference between the informative and uninformative conditions, at the group level (N = 44).

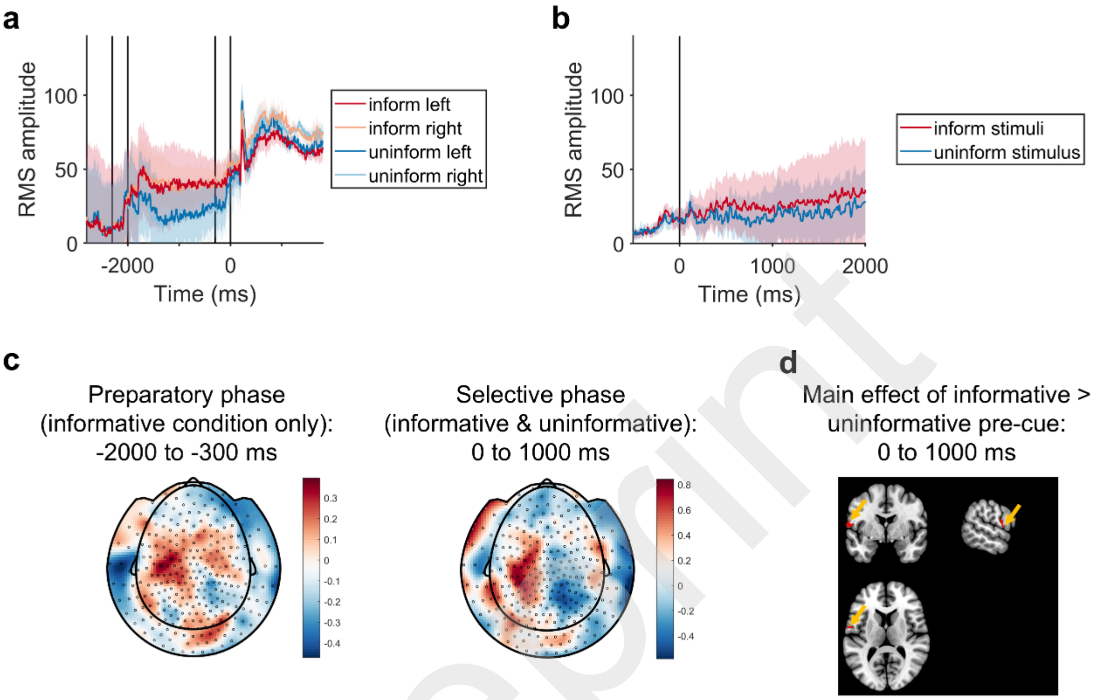


Preprint

**Supplementary Fig. 2. (continued on next page)** Results from the source analysis in 100-ms time windows throughout the preparatory phase (between -1800 and -300 ms). Each image shows the difference between the informative and uninformative conditions at the group level (N = 44). The left column shows glass brain images, and the other columns show the functional activity superimposed onto structural images, centered on co-ordinates in the visual cortex ([-4, 92, 6]), left auditory cortex ([-56, 0, 2]), right inferior frontal gyrus (IFG; [54, 18, 26]), and left superior parietal lobule (SPL; [-22, -42, 54]).

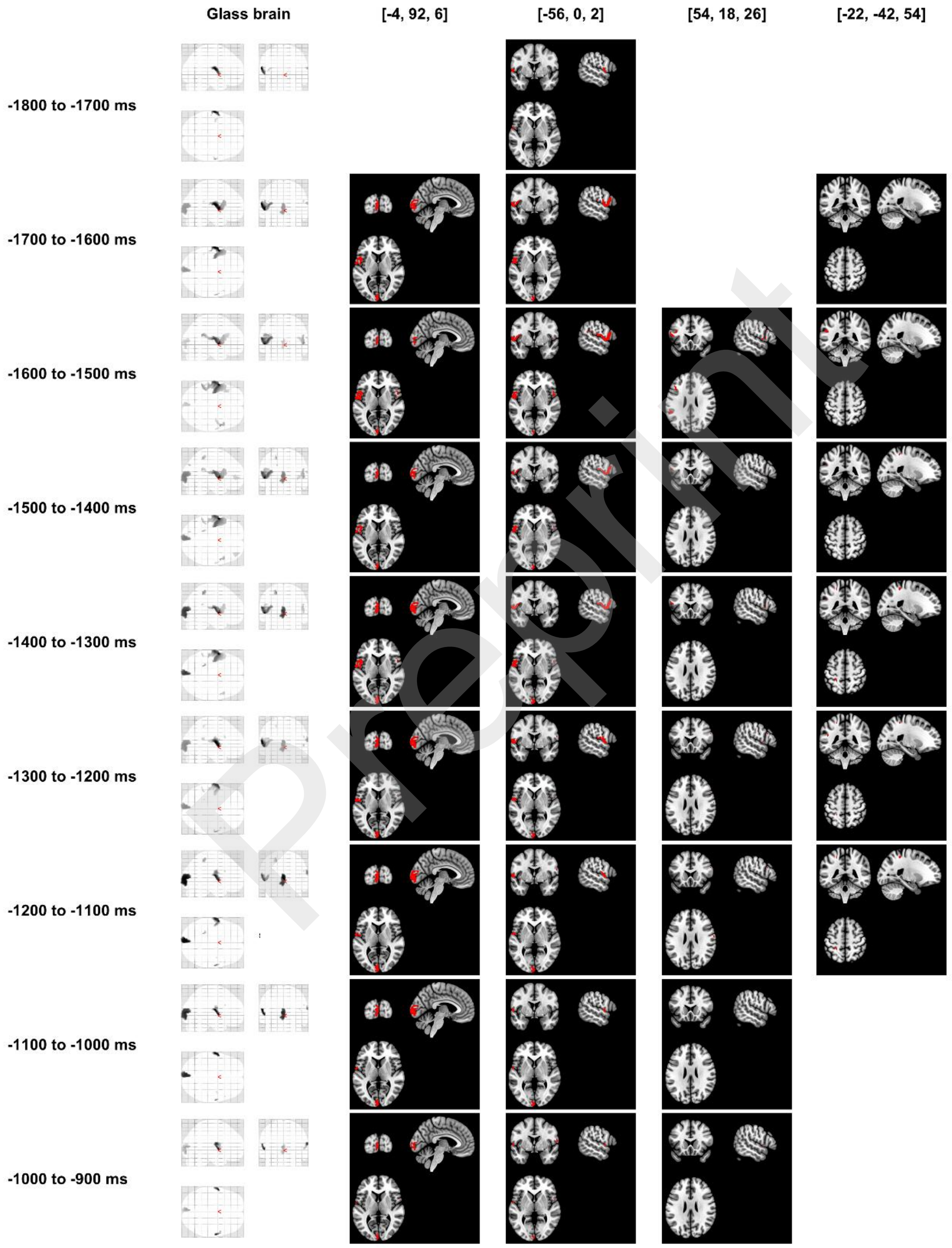


Preprint

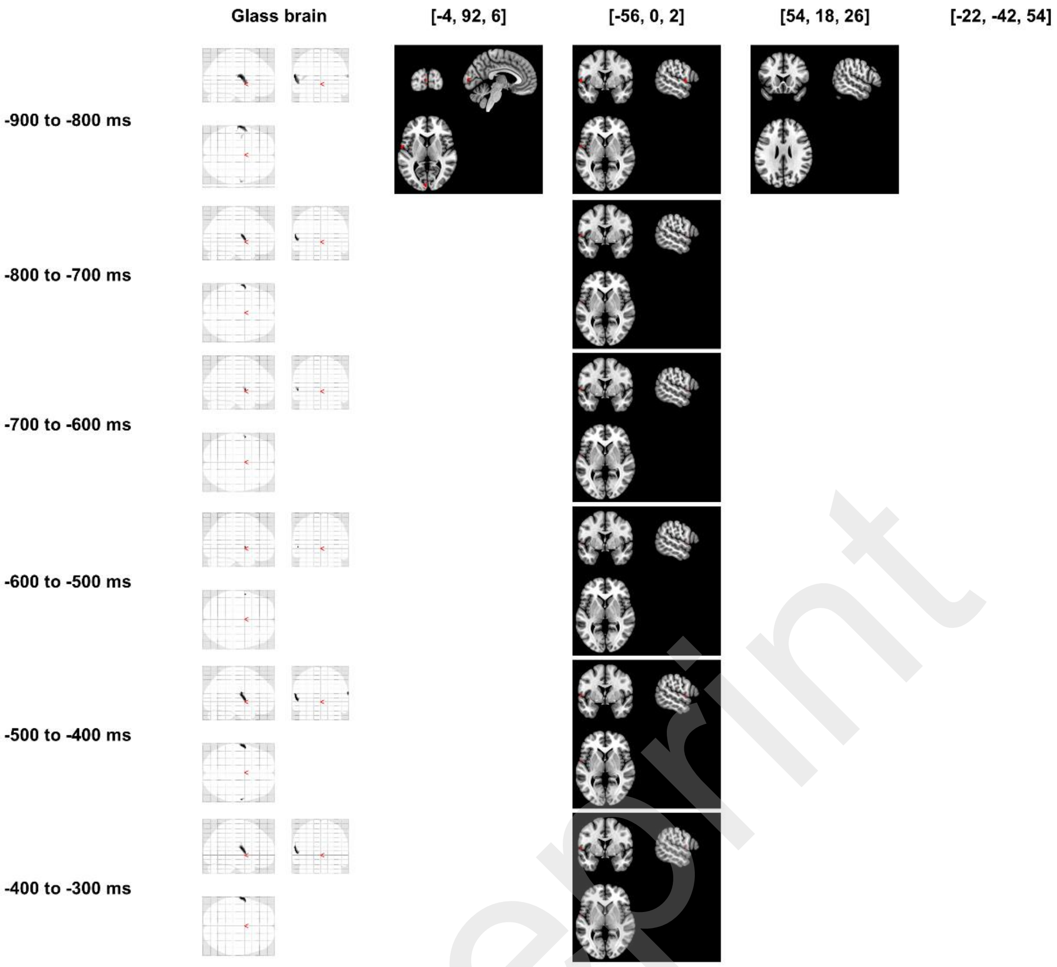
Glass brain
[-4, 92, 6]
[-56, 0, 2]
[54, 18, 26]
[-22, -42, 54]
-900 to -800 ms
-800 to -700 ms
-700 to -600 ms
-600 to -500 ms
-500 to -400 ms
-400 to -300 ms

Preprint

**Supplementary Fig. 3.** Pupil diameter across the group, following foreshortening error correction in preprocessing. Lines show the means and standard errors in the informative and uninformative conditions, across participants who had usable pupil data (N = 30).

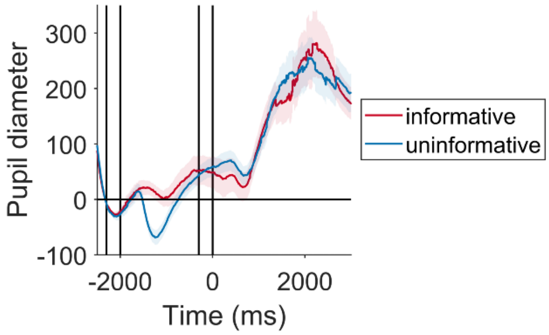


Preprint

**Supplementary Table 1.** Results of MEG source analysis for the preparatory phase. Coordinates as [x, y, z] in MNI space. Areas were labelled using the Maximum Probability Map from the JuBrain Anatomy Toolbox for SPM (1) (version 3.0); percentages reflect the percentage of the cluster volume in each area.

| *Peak co-ordinates (mm)* | *Area(s)* | *Number of voxels* | $p_{FWE}$ |
|---|---|---|---|
| [-62, -8, 14] | Area OP4 [PV] (21.3%)<br>Area TE 1.2 (4.6%)<br>Area TE 3 (4.0%)<br>Area OP1 [SII] (2.8%)<br>Area 3a (0.8%)<br>Area TE 4 (0.4%)<br>Area 3b (0.2%) | 226 | .021 |
| [-6, -92, 6] | Area hOc1 [V1] (58.0%)<br>Area hOc2 [V2] (27.4%) | 336 | .016 |
| [64, -10, 18] | Area OP4 [PV] (59.2%)<br>Area 3b (25.0%)<br>Area PFop (IPL) (5.4%)<br>Area OP1 [SII] (1.1%) | 23 | .041 |
| [-50, -44, 20] | Area PFcm (IPL) (5.1%) | 17 | .043 |
| [-54, -46, 20] | Area PFm (IPL) (4.9%)<br>Area PGa (IPL) (2.1%) | 1 | .049 |

Preprint

**Supplementary Table 2.** Results of MEG source analysis for sub-windows within the preparatory phase, for the group-level comparison between the informative and uninformative conditions. Time windows with no significant sources are not shown. Areas were labelled using the Maximum Probability Map from the JuBrain Anatomy Toolbox for SPM (1) (version 3.0); percentages reflect the percentage of the cluster volume in each area. For clusters in which no percentages are provided for area labels, the Maximum Probability Map returned no results, and area labels instead reflect those with a greater than zero probability at the cluster location.

| *Time window (ms)* | *Peak co-ordinates (mm)* | *Area(s)* | *Number of voxels* | $p_{FWE}$ |
|---|---|---|---|---|
| -1800 to -1700 | [-64, -12, 16] | Area OP4 [PV] (26.6%)<br>Area OP1 [SII] (8.3%)<br>Area TE 1.2 (6.2%)<br>Area TE 3 (4.3%)<br>Area 3b (0.1%)<br>Area TE 4 (0.1%) | 152 | .024 |
| | [64, -10, 16] | Area OP4 [PV] (77.3%)<br>Area 3b (11.7%)<br>Area OP1 [SII] (10.2%) | 16 | .042 |
| | [-54, -20, 18] | Area OP1 [SII] (100.0%) | 2 | .048 |
| | [-44, -8, 12] | Area OP3 [VS] (29.8%)<br>Area Ig3 (23.1%)<br>Area Id4 (10.6%)<br>Area Id5 (9.6%)<br>Area OP4 [PV] (1.9%)<br>Area Id2 (1.0%) | 13 | .043 |
| | [-46, -4, 8] | Area OP4 [PV]<br>Area Id5<br>Area Id4<br>Area Id2<br>Area TE 1.2<br>Area Id6 | 1 | .049 |
| | [58, -14, 18] | Area OP1 [SII] (100.0%) | 1 | .049 |
| -1700 to -1600 | [-62, -8, 14] | Area 44 (16.1%)<br>Area OP4 [PV] (8.0%)<br>Area OP1 [SII] (3.6%)<br>Area TE 1.2 (2.5%)<br>Area OP3 [VS] (2.3%)<br>Area Id4 (2.0%)<br>Area OP8 (1.6%)<br>Area Ig3 (1.5%)<br>Area TE 3 (1.1%)<br>Area Id5 (0.8%)<br>Area Id2 (0.3%)<br>Area 3a (0.3%)<br>Area 45 (0.2%)<br>Area TE 4 (0.1%)<br>Area OP9 (0.1%)<br>Area 3b (0.1%) | 676 | .007 |
| | [-2, -90, -6] | Area hOc1 [V1] (49.2%)<br>Area hOc2 [V2] (34.8%) | 320 | .015 |
| | [62, -4, 18] | Area OP4 [PV] (69.6%)<br>Area 3b (27.7%)<br>Area OP1 [SII] (1.8%) | 14 | .043 |
| | [52, 14, 22] | Area 44 (75.0%) | 5 | .046 |
| | [60, 0, 18] | Area 3b<br>Area 3a<br>Area OP4 [PV]<br>Area 4p | 2 | .048 |
| | [-44, -40, 22] | Area PFcm (IPL) (25.0%) | 2 | .048 |

Preprint

| | | | | |
|---|---|---|---|---|
| -1600 to -1500 | [-56, 0, 0] | Area 44 (12.4%)<br>Area OP4 [PV] (10.1%)<br>Area OP1 [SII] (7.8%)<br>Area PFcm (IPL) (5.2%)<br>Area TE 1.2 (3.2%)<br>Area OP3 [VS] (2.8%)<br>Area Id4 (2.8%)<br>Area Ig3 (2.0%)<br>Area OP8 (1.9%)<br>Area TE 1.0 (1.5%)<br>Area TE 3 (0.9%)<br>Area Id5 (0.7%)<br>Area Id2 (0.5%)<br>Area TE 1.1 (0.5%)<br>Area TE 4 (0.2%)<br>Area 45 (0.1%)<br>Area OP9 (0.1%)<br>Area PFop (IPL) (0.1%) | 910 | .002 |
| | [54, 6, 2] | Area 44 (40.5%)<br>Area OP3 [VS] (4.6%)<br>Area OP8 (0.5%)<br>Area Id4 (0.4%)<br>Area TE 3 (0.3%)<br>Area TE 1.2 (0.1%) | 149 | .027 |
| | [-2, -82, 8] | Area hOc1 [V1] (58.5%)<br>Area hOc2 [V2] (25.2%) | 145 | .027 |
| | [-38, 20, 20] | Area 44<br>Area 45 | 65 | .035 |
| | [62, -4, 18] | Area OP4 [PV] (50.0%)<br>Area 3b (47.9%) | 6 | .046 |
| | [60, 0, 18] | Area 3b<br>Area 3a<br>Area OP4 [PV]<br>Area 4p | 2 | .048 |
| -1500 to -1400 | [-56, 2, 0] | Area 44 (15.0%)<br>Area OP4 [PV] (10.5%)<br>Area OP1 [SII] (5.3%)<br>Area TE 1.2 (3.5%)<br>Area Id4 (3.2%)<br>Area OP3 [VS] (3.1%)<br>Area OP8 (3.0%)<br>Area Ig3 (2.0%)<br>Area TE 1.0 (1.6%)<br>Area TE 3 (1.1%)<br>Area Id5 (0.6%)<br>Area Id2 (0.6%)<br>Area 45 (0.5%)<br>Area TE 4 (0.2%)<br>Area TE 1.1 (0.1%)<br>Area OP9 (0.1%)<br>Area Id6 (0.1%) | 903 | .008 |
| | [-4, -92, 6] | Area hOc1 [V1] (48.2%)<br>Area hOc2 [V2] (34.7%)<br>Area hOc3d [V3d] (0.1%) | 393 | .017 |
| | [52, 6, 2] | Area 44 (59.1%)<br>Area OP8 (0.7%)<br>Area TE 3 (0.4%)<br>Area TE 1.2 (0.1%) | 103 | .032 |
| | [-22, -42, 54] | Area 5L (SPL) (36.2%)<br>Area 2 (4.7%) | 29 | .041 |

Preprint

| | | | | |
|---|---|---|---|---|
| | | Area 3b (0.9%) | | |
| | [-52, -44, 24] | Area PFcm (IPL) (38.9%)<br>Area TE 1.1 (2.5%)<br>Area OP1 [SII] (0.8%) | 45 | .039 |
| | [-20, -80, 34] | Area hIP7 (IPS) (62.5%)<br>Area hPO1 (IPS) (27.5%)<br>Area hOc4d [V3A] (10.0%) | 5 | .047 |
| | [36, 48, -2] | Area Fo5 (2.2%) | 46 | .039 |
| | [62, -4, 16] | Area OP4 [PV] (74.0%)<br>Area 3b (23.1%) | 13 | .045 |
| | [60, 0, 18] | Area 3b<br>Area OP4 [PV]<br>Area 3a<br>Area 44<br>Area 4p | 3 | .048 |
| -1400 to -1300 | [-56, 0, 0] | Area 44 (12.0%)<br>Area OP4 [PV] (11.7%)<br>Area OP1 [SII] (4.6%)<br>Area TE 1.2 (4.2%)<br>Area Id4 (3.5%)<br>Area OP3 [VS] (2.9%)<br>Area Ig3 (2.4%)<br>Area OP8 (1.8%)<br>Area TE 1.0 (1.6%)<br>Area TE 3 (1.3%)<br>Area Id5 (0.8%)<br>Area Id2 (0.5%)<br>Area TE 4 (0.2%)<br>Area OP9 (0.1%) | 721 | .010 |
| | [-4, -92, 6] | Area hOc1 [V1] (48.1%)<br>Area hOc2 [V2] (34.4%)<br>Area hOc3d [V3d] (0.1%) | 404 | .017 |
| | [52, 12, 14] | Area 44 (68.9%) | 68 | .036 |
| | [-22, -42, 54] | Area 5L (SPL) (33.9%)<br>Area 2 (14.5%)<br>Area 3b (0.7%) | 38 | .040 |
| | [-52, -44, 24] | Area PFcm (IPL) (16.7%) | 9 | .046 |
| -1300 to -1200 | [-56, 0, 2] | Area OP4 [PV] (20.0%)<br>Area OP1 [SII] (7.2%)<br>Area TE 1.2 (6.6%)<br>Area OP3 [VS] (3.0%)<br>Area TE 3 (2.4%)<br>Area Ig3 (2.2%)<br>Area TE 1.0 (1.8%)<br>Area Id4 (1.7%)<br>Area Id5 (1.3%)<br>Area Id2 (0.9%)<br>Area 3a (0.4%)<br>Area TE 4 (0.2%)<br>Area 3b (0.1%) | 436 | .011 |
| | [-2, -90, -6] | Area hOc1 [V1] (47.0%)<br>Area hOc2 [V2] (35.6%) | 398 | .012 |
| | [64, -8, 18] | Area OP4 [PV] (46.2%)<br>Area 3b (10.3%)<br>Area OP1 [SII] (7.1%)<br>Area PFop (IPL) (3.2%) | 62 | .033 |
| | [54, 6, 2] | Area OP4 [PV]<br>Area 44 | 7 | .045 |

Preprint

| | | | | |
|---|---|---|---|---|
| | | Area OP8<br>Area TE 1.2<br>Area TE 3 | | |
| | [-44, -42, 22] | Area PFcm (IPL) (4.4%) | 20 | .041 |
| | [-22, -42, 54] | Area 5L (SPL) (18.8%) | 4 | .047 |
| | [54, 18, 24] | Area 44 (71.9%)<br>Area 45 (15.6%) | 4 | .047 |
| | [-24, -38, 58] | Area 5L (SPL) (100.0%) | 2 | .048 |
| -1200 to -1100 | [-2, -90, -6] | Area hOc1 [V1] (47.7%)<br>Area hOc2 [V2] (35.3%)<br>Area hOc3d [V3d] (0.1%) | 433 | .012 |
| | [-56, 0, 2] | Area OP4 [PV] (27.4%)<br>Area TE 1.2 (12.2%)<br>Area TE 3 (4.3%)<br>Area Ig3 (1.6%)<br>Area Id5 (1.3%)<br>Area OP1 [SII] (1.1%)<br>Area Id2 (1.0%)<br>Area OP3 [VS] (0.8%)<br>Area TE 4 (0.4%)<br>Area 3b (0.2%) | 214 | .021 |
| | [62, -6, 22] | Area OP4 [PV] (30.3%)<br>Area 3b (26.8%)<br>Area PFop (IPL) (3.3%)<br>Area OP1 [SII] (2.0%) | 57 | .035 |
| | [-22, -42, 54] | Area 5L (SPL) (29.4%)<br>Area 2 (13.2%)<br>Area 3b (0.7%) | 34 | .039 |
| | [54, 18, 24] | Area 44 (79.2%)<br>Area 45 (20.8%) | 3 | .048 |
| | [54, 6, 2] | Area OP4 [PV]<br>Area TE 3<br>Area 44<br>Area OP8<br>Area TE 1.2 | 1 | .049 |
| | [42, -6, 16] | Area OP3 [VS] (62.5%) | 1 | .049 |
| -1100 to -1000 | [-56, 0, 2] | Area OP4 [PV] (38.6%)<br>Area TE 1.2 (9.4%)<br>Area TE 3 (8.1%)<br>Area OP1 [SII] (2.4%)<br>Area TE 4 (0.9%)<br>Area 3b (0.1%) | 94 | .029 |
| | [-6, -92, 6] | Area hOc1 [V1] (48.7%)<br>Area hOc2 [V2] (33.9%) | 366 | .013 |
| | [64, -8, 18] | Area OP4 [PV] (56.9%)<br>Area 3b (31.9%) | 20 | .041 |
| | [42, -6, 16] | Area OP3 [VS] (62.5%) | 1 | .049 |
| | [54, 18, 26] | Area 45 (62.5%)<br>Area 44 (37.5%) | 1 | .049 |
| | [46, -4, 12] | Area OP3 [VS]<br>Area OP4 [PV]<br>Area Id4 | 1 | .049 |
| -1000 to -900 | [-56, 0, 2] | Area OP4 [PV] (35.9%)<br>Area TE 1.2 (13.0%)<br>Area TE 3 (11.2%)<br>Area TE 4 (0.3%) | 48 | .036 |
| | [62, -6, 16] | Area OP4 [PV] (33.1%)<br>Area 3b (4.7%)<br>Area TE 3 (0.3%) | 74 | .033 |

Preprint

| | | | | |
|---|---|---|---|---|
| | [-6, -92, 6] | Area hOc1 [V1] (57.3%)<br>Area hOc2 [V2] (27.3%) | 162 | .024 |
| -900 to -800 | [-56, 0, 2] | Area OP4 [PV] (27.8%)<br>Area TE 1.2 (8.6%)<br>Area TE 3 (6.9%)<br>Area 3b (0.2%)<br>Area TE 4 (0.2%) | 80 | .032 |
| | [-2, -82, 8] | Area hOc1 [V1] (77.2%)<br>Area hOc2 [V2] (20.3%) | 40 | .038 |
| | [64, -10, 20] | Area OP4 [PV] (70.0%)<br>Area PFop (IPL) (8.3%)<br>Area 3b (6.7%)<br>Area OP1 [SII] (2.5%) | 15 | .043 |
| | [-50, -22, 14] | Area OP1 [SII] (48.4%)<br>Area TE 1.0 (45.3%) | 8 | .045 |
| | [50, 16, 30] | Area 45 (27.5%)<br>Area 44 (22.5%) | 5 | .047 |
| -800 to -700 | [-60, 2, 6] | Area OP4 [PV] (29.2%)<br>Area TE 3 (11.4%)<br>Area TE 1.2 (8.3%)<br>Area 3b (0.4%) | 33 | .039 |
| -700 to -600 | [-56, 0, 2] | Area TE 3 (25%)<br>Area TE 1.2 (25%)<br>Area OP4 [PV] (1.1%) | 11 | .045 |
| -600 to -500 | [-56, 0, 2] | Area TE 3 (6.2%) | 2 | .048 |
| -500 to -400 | [-56, 0, 2] | Area OP4 [PV] (40.1%)<br>Area TE 1.2 (12.9%)<br>Area TE 3 (9.6%)<br>Area 3b (0.2%)<br>Area TE 4 (0.2%) | 57 | .035 |
| | [64, -10, 18] | Area OP4 [PV] (77.8%)<br>Area 3b (22.2%) | 9 | .045 |
| -400 to -300 | [-62, -8, 12] | Area OP4 [PV] (36.8%)<br>Area TE 1.2 (7.9%)<br>Area TE 3 (6.2%)<br>Area OP1 [SII] (0.9%)<br>Area 3b (0.2%) | 68 | .034 |
| | [-62, -10, 24] | Area OP4 [PV] (36.8%)<br>Area TE 1.2 (7.9%)<br>Area TE 3 (6.2%)<br>Area OP1 [SII] (0.9%)<br>Area 3b (0.2%) | 2 | .048 |

Preprint

**Supplementary Table 3.** Results of MEG source analysis for sub-windows within the preparatory phase, for the between-subjects covariates. Coordinates as [x, y, z] in MNI space. Time windows without significant sources are not shown. Areas were labelled using the Maximum Probability Map from the JuBrain Anatomy Toolbox for SPM (1) (version 3.0); percentages reflect the percentage of the cluster volume in each area. For clusters in which no percentages are provided for area labels, the Maximum Probability Map returned no results, and area labels instead reflect those with a greater than zero probability at the cluster location.

| **Interaction between informative versus uninformative and audiometric thresholds** | | | | |
|---|---|---|---|---|
| *Time window (ms)* | *Peak co-ordinates (mm)* | *Area(s)* | *Number of voxels* | $p_{FWE}$ |
| -1400 to -1300 | [-28, -30, 52] | Area 3a (82.5%)<br>Area 4p (17.5%) | 5 | .047 |
| -800 to -700 | [-26, -30, 54] | Area 3a (52.7%)<br>Area 4p (9.3%)<br>Area 3b (4.5%) | 47 | .036 |

Preprint

**Supplementary Table 4.** Results of MEG source analysis for the selective phase. Coordinates as [x, y, z] in MNI space. Areas were labelled using the Maximum Probability Map from the JuBrain Anatomy Toolbox for SPM (1) (version 3.0); percentages reflect the percentage of the cluster volume in each area.

| *Peak co-ordinates (mm)* | *Area(s)* | *Number of voxels* | $p_{FWE}$ |
|---|---|---|---|
| [-58, -2, 10] | Area OP4 [PV] (5.6%)<br>Area 3a (1.5%)<br>Area TE 3 (1.1%)<br>Area TE 1.2 (1.0%)<br>Area 3b (0.4%) | 125 | .031 |

Preprint

**Supplementary Table 5**. Results of MEG source analysis for sub-windows within the preparatory phase, for the pupil covariates. Coordinates as [x, y, z] in MNI space. Time windows without significant sources are not shown. Areas were labelled using the Maximum Probability Map from the JuBrain Anatomy Toolbox for SPM (1) (version 3.0); percentages reflect the percentage of the cluster volume in each area. For clusters in which no percentages are provided for area labels, the Maximum Probability Map returned no results, and area labels instead reflect those with a greater than zero probability at the cluster location.

| **Interaction between informative versus uninformative and Preparatory Phase pupil diameter squared** | | | | |
|---|---|---|---|---|
| *Time window (ms)* | *Peak co-ordinates (mm)* | *Area(s)* | *Number of voxels* | $p_{FWE}$ |
| -1900 to -1800 | [-32, 44, 28] | Area 45<br>Area OP9 | 112 | .024 |
| | [-6, 48, 38] | Area p32 | 5 | .046 |
| -1800 to -1700 | [-32, 44, 28] | Area 45<br>Area OP9<br>Area Id7 | 289 | .013 |
| | [-6, 50, 34] | Area p32 (38.4%) | 101 | .025 |
| -1700 to -1600 | [-32, 44, 28] | Area 45<br>Area OP9 | 138 | .021 |
| -1600 to -1500 | [-36, 42, 26] | Area 45<br>Area OP9<br>Area Id7 | 183 | .022 |
| -1500 to -1400 | [-42, 38, 18] | Area 45<br>Area OP9<br>Area Id7<br>Area OP8 | 238 | .026 |
| -1400 to -1300 | [-42, 38, 18] | Area 45<br>Area OP9<br>Area Id7 | 225 | .029 |
| | [42, 40, 12] | Area 45 | 9 | .047 |
| | [42, 34, 16] | Area 45 | 2 | .049 |

Preprint

### References

1. S. B. Eickhoff, *et al.*, A new SPM toolbox for combining probabilistic cytoarchitectonic maps and functional imaging data. *Neuroimage* **25**, 1325–1335 (2005).

Preprint
